# Self-thresholding hierarchical outlier-detection for animal movement tracks

**DOI:** 10.64898/2026.07.11.737894

**Authors:** Kamran Safi

**Affiliations:** Max Planck Institute of Animal Behavior, Radolfzell, Germany; Department of Biology, University of Konstanz, Konstanz, Germany

**Keywords:** animal movement, Brownian bridge, GPS spoofing, GPS telemetry, location error, move2, outlier detection, quality control

## Abstract

Erroneous locations are ubiquitous in animal tracking, propagating into estimates of movement rate, home range, habitat selection and behavioural state. Existing filters often need a threshold chosen per species and schedule, assume a movement process heterogeneous data rarely satisfy, or stop at a plot for manual filtering. A block of consecutive wrong positions, caused for example by GNSS spoofing, whose interior looks ordinary step by step, is caught only by one supervised filter.

I introduce mt_clean_track(), an unsupervised cleaning cascade in the R package move2utils. Four detectors score every location, each for an error class the others cannot: a Brownian-bridge geometric residual, a movement- probability score, a path-versus-displacement detour ratio, and a speed cap. A fifth stage, not a detector, identifies a block of false positions at a track’s start or end, where no per-location test can tell which side is the artefact. Every threshold bar the detour ratio’s is read from the track’s own scores, and a location is flagged only on corroborated evidence, so the composition controls the false-positive rate.

I evaluate the cascade on synthetic trajectories with known truth, spanning spikes, spoofing blocks, colony halos and multi-state anomalies, against four alternatives (a naïve speed cap, atlastools, trip::sda, SDLfilter). Each alternative is given the same speed ceiling, calibrated on the cohort’s uncontaminated track; the cascade, nothing but the data. Even so, it recovered all 141 injected outliers on every contaminated track, whereas they left 27 to 33 behind at recalls of 0.766–0.809. No alternative scored better on recall and precision together; where it ranked below one, the cost was -precision, never a retained outlier. On a coherent spoofing block it recovered 30 of 30 injected locations against their 0 to 3; on a clean track with 30 m positional error it flagged one location across 100 realisations.

Unsupervised operation, data-driven thresholds and a read-out of what triggered each flag make outlier screening routine and reproducible instead of bespoke. The cascade screens rather than models, and the functions beneath it stay open to further detectors and to new consensus-finding rules.

## 1 Introduction

Animals’ paths encode information about their physiology, behaviour, and the ways they interact with the landscapes they inhabit and traverse. Tracking technologies record these paths at various resolutions, generating millions of locations annually across hundreds of species and thousands of individuals, feeding into analyses of space use, resource selection, energetics, and social behaviour (Kays *et al*., 2022). Location estimates, like all measurements, carry error (Péron *et al*., 2020): GPS locations can be displaced by tens to hundreds of metres through poor satellite geometry, signal multipath, or obstruction (Frair *et al*., 2010; Lewis *et al*., 2007; D’Eon & Delparte, 2005). A more recent and increasingly pervasive source of error is deliberate GNSS jamming and spoofing in conflict zones, where military countermeasures feed false signals to GPS receivers (Jiguet *et al*., 2025; Martín-Vélez *et al*., 2026). Whatever the source, erroneous locations distort down- stream estimates of speed, turning-angle distributions, home-range size, habitat-selection coefficients, and behavioural-state assignments (Hurford, 2009; Bjørneraas *et al*., 2010; Gupte *et al*., 2022; Frair *et al*., 2010). Errors can reach a magnitude where we would consider them outliers, which are not a different kind of location, only locations far enough out that keeping them would distort those said estimates. That boundary is a convention, as it is wherever statistics, or subjective decisions of experts, separate a tail from a bulk. Objectively, we never really know that the locations we keep are accurate, only which are too far out to keep.

Screening and removing outliers is one of the first and consequential steps in any movement-analysis pipeline, yet the methods in use are largely bespoke to a given species, device, and sampling schedule, adding manual work and lowering reproducibility. The simplest and most widely used methods are threshold filters: a maximum plausible speed (McConnell *et al*., 1992), a speed–distance–angle criterion (Freitas *et al*., 2008; Austin *et al*., 2003), or turning-angle rules (Douglas *et al*., 2012; Shimada *et al*., 2012; Bjørneraas *et al*., 2010). While computationally cheap, they demand species and sampling schedule specific parameters, return binary retain-or-remove decisions with no probabilistic gradation, and they are scale-sensitive: the same species needs different thresholds at different sampling intervals, so locations are misclassified whenever intervals vary and/or behaviour shifts.

State-space and continuous-time movement models instead treat locations as noisy observations of a latent process and return smoothed estimates (Jonsen *et al*., 2005; Johnson *et al*., 2008; Jonsen *et al*., 2023); they are statistically principled but computationally demanding, and each commits to a specific movement process that real data need not follow. Diagnostic tools such as ctmm::outlie() (Calabrese *et al*., 2016; Fleming *et al*., 2020) rely on an assumed measurement-error (UERE) derived from calibration and visualise speed and distance anomalies but leave the removal decision to the analyst.

Existing approaches share yet a deeper limitation. Filters and smoothers, whether parametric or empirical, reduce each location to a single judgement about its transition to neighbours: is the speed, the turning angle plausible, is the residual to the smoothed path plausible? Some consequential outliers in real tracking data, however, come in *spatially coherent blocks*: a GPS receiver locks onto a reflected signal and produces a drift cluster of positions that sit together in the wrong place, or a spoofing/jamming event pins the receiver at a single falsified coordinate for minutes to hours (Jiguet *et al*., 2025; Martín-Vélez *et al*., 2026). The eye sees such clusters at once as they are geometrically detached from where the animal should be, whether or not within-block transition is plausible. Because such blocks have normal interiors and extreme boundaries, a test built on one-step transitions can detect the entry and exit of the block; it is however structurally blind to the interior. Comparing a location with a *window* of its neighbours escapes that blindness (Gupte *et al*., 2022). Although a first attempt exists in the package atlastools, the challenge remains open to an unsupervised treatment.

I propose an approach which combines by design complementary signals, such that

i. each location receives a score reflecting how unusual its movement is under the animal’s own empirical distribution of transitions, (ii) each location also receives a score reflecting how far it deviates from where its temporal neighbours place it geometrically,
ii. thresholds are determined automatically from the empirical structure of the score distribution rather than from user-supplied parameters, (iv) irregular sampling is handled by construction rather than as an afterthought, and (v) a block of consecutive outliers is treated as one object, so that the gap above is addressed rather than noted.

Here, I introduce these detectors as major components of the move2utils R package for working with move2 objects (Kranstauber *et al*., 2024); the package as a whole is described in a companion Application Note (Kranstauber *et al*., 2026). They build on established machinery: the empirical step-and-turn architecture introduced for trajectory simulation by Technitis (2021), the Brownian bridge (Kranstauber *et al*., 2012) and its directional variant (Kranstauber *et al*., 2014), and self-determining thresholds (MacArthur,1957; Silverman, 1981).

## 2 A cleaning cascade for movement tracks

The function mt_clean_track() takes a movement track (a sequence of timestamped GPS positions packaged as a move2 object (Kranstauber *et al*., 2024)) and by default returns it cleaned.

Within mt_clean_track(), several detectors are arranged as a cascade to operate in a specific order, each looking at the same track, yet none of them deciding on its own. The cascade the function runs through rests on four pillars: first, four detectors that fail in different ways; second, a threshold each detector reads off the track it is given; third, a consensus-finding rule that requires corroboration; and fourth, a segmentation step that recovers blocks of outliers no detector can see. Ahead of all four, when a physiological ceiling is supplied, the cascade peels away the endpoints of steps no animal could have made (Figure 1a). Supplement S1 gives every formulation.

**Figure 1:**
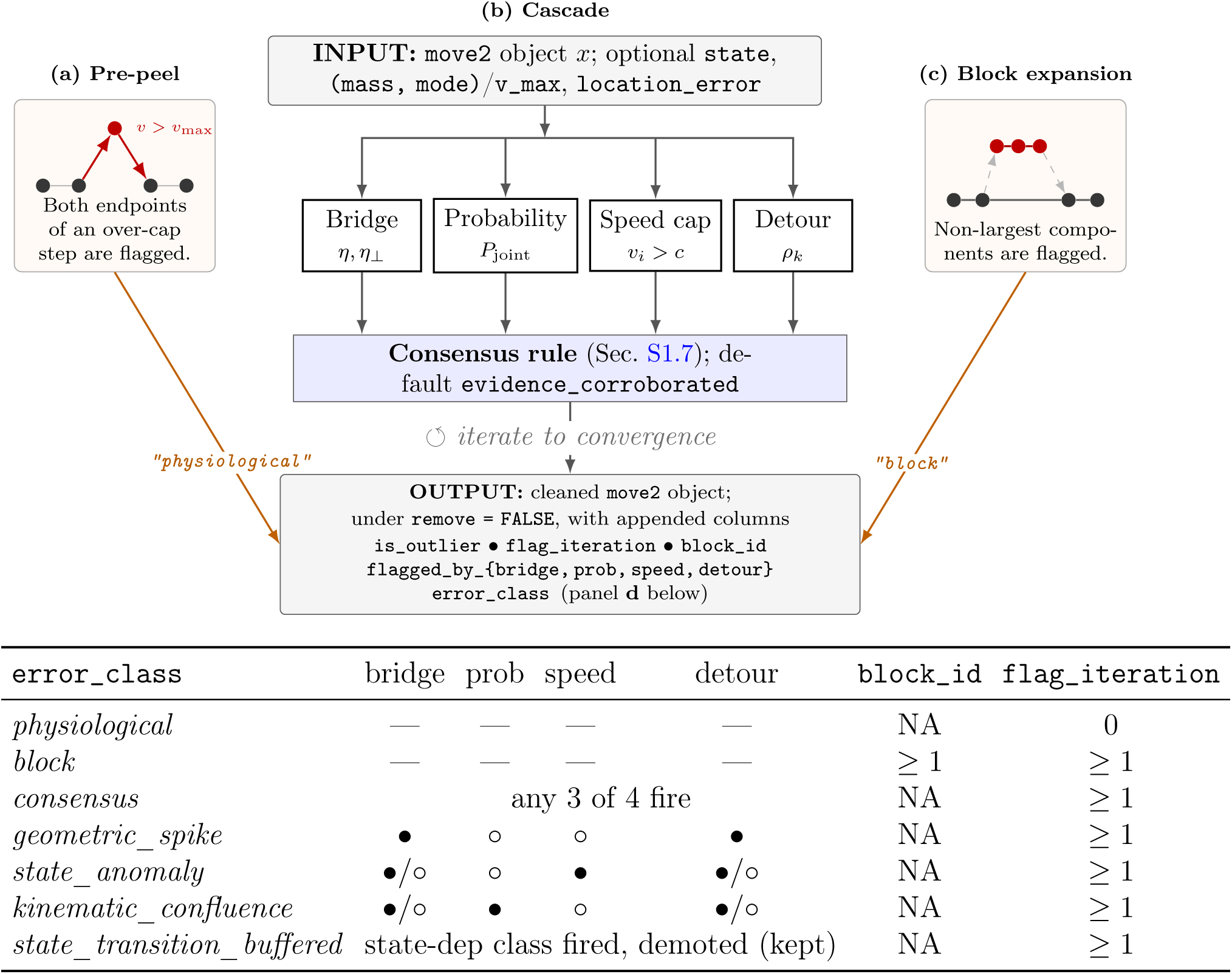
The. mt_clean_track() **cleaning cascade. (a) Pre-peel (left blow-out):** when a physiological speed bound is supplied, iterative pre-peeling flags both endpoints of every over-cap step before any per-location detector runs; these carry error_class = "physiological". **(b) Cascade (centre):** the four self-thresholding detectors feed the consensus-finding rule (default evidence_corroborated); ⟲ marks iteration to conver- gence. **(c) Block expansion (right blow-out):** segments the kept locations along the track and flags every location outside the largest segment as error_class = "block". (a) **Output fingerprint by** error_class: filled circle ( ) the detector fired, open cir- cle ( ) it was silent, “ */* ” either (the rule does not constrain it), “—” not part of the rule. error_class reports the highest-priority class that fired, top to bottom, while the flagged_by_* columns keep the detector-level firings. The predicates shown are those of the class_aware rule; the default evidence_corroborated decides by accumulating evidence, so a location can be flagged without satisfying any predicate here and is then reported unclassified. Cross-track pooling adds a "pool" class not shown (§ 4).

A location is flagged only when the four detectors corroborate one another based on a chosen consensus-finding rule. What each detector brings to the decision is not the size of its own score, but how far past its own threshold a location lies. A detector whose numbers run large therefore cannot dominate one whose numbers run small. The function runs through the cascade repeatedly, re-fitting each threshold on the locations that survived, until a pass adds no new flags or the limit is reached.

Thus, mt_clean_track() decides which locations are outliers and by default drops them; asked to keep them with remove = FALSE, it marks each one and attaches the evidence behind each decision. The function has thirty-three arguments but requires exactly one, the track. The rest are knobs and levers, none of which have to be turned. Everything it uses, it reads off the track it is given, and unless a table says otherwise every result in this paper comes from that single-argument call. What comes back is a move2 object of the same form as the input, ready for home-range estimation, step-selection models or behavioural segmentation.

### The four detectors

No single detector is suited for every source of error. Real tracks carry at least four broad classes of contamination: geometric anomalies, where a location sits implausibly far from where its neighbours place it; kinematic anomalies, where the speeds and turns it implies are unlike the animal’s own; violations of a biological speed bound, and symmetric out-and-back excursions, which have ordinary kinematics and pathological geometry. The cascade runs one complementary detector per class (*bridge*, *probability*, *speed cap* and *detour* ). Table 1 sets out what each catches and where it fails.

**Table 1:**
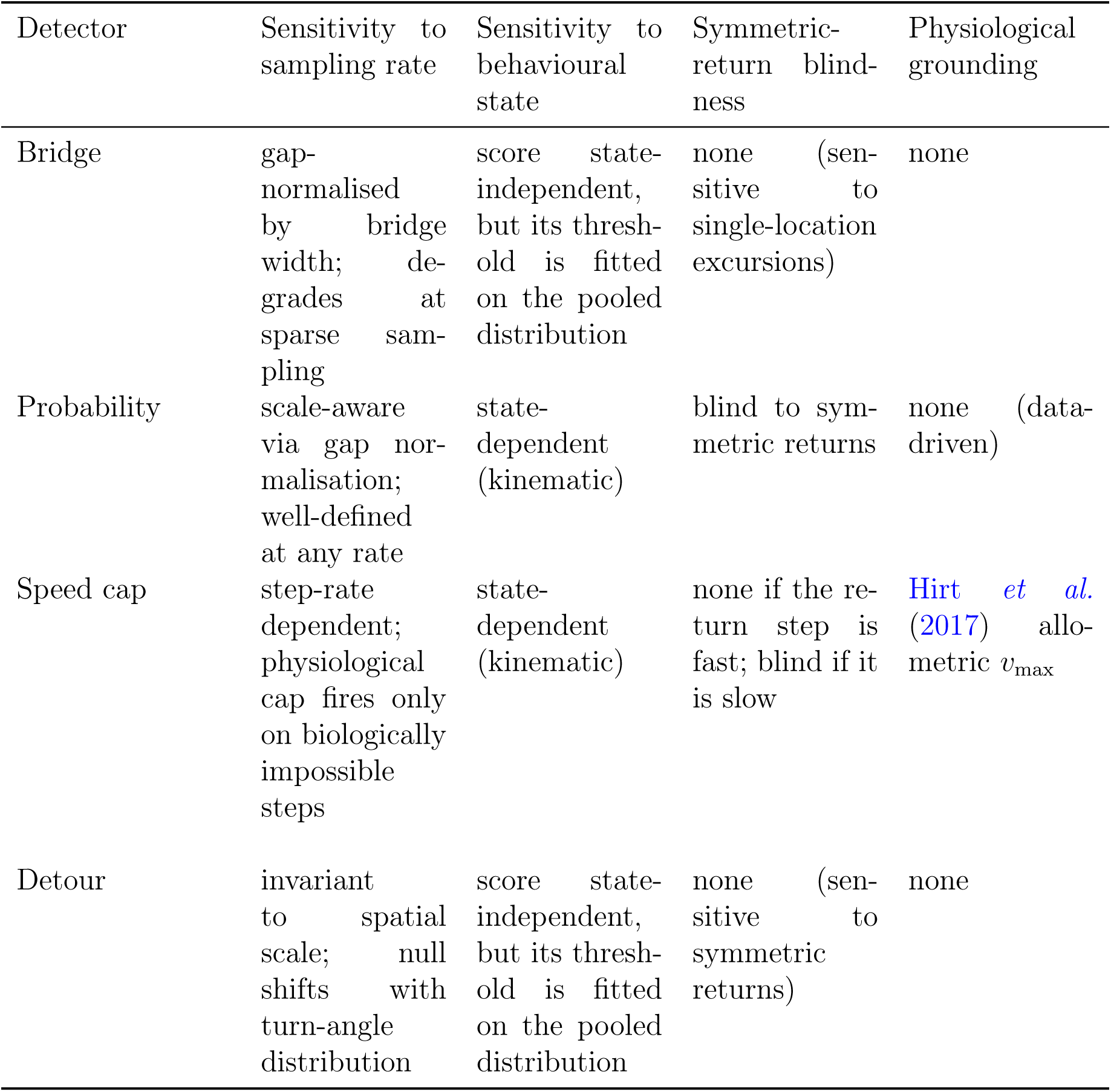
Scope of validity for each detector, named as in the text. Sensitivity to sampling rate is the detector’s response to a uniform increase in the interval between locations; sensitivity to behavioural state, to mixing distinct activity distributions in one track; symmetric-return blindness, to a colony-halo or out-and-back spike at modest displace- ment. Physiological grounding is whether the detector uses an external biological prior <u>(mass, mode).</u>

### The bridge detector

measures how far a location lies from where its neighbours place it, divided by a width built only from the timestamps. That expected position and width are those of a Brownian bridge, the diffusion path pinned at both neighbours: the mean is their time-weighted midpoint and the width the spread such a path would show at that moment. Because that width uses no positions, an outlier cannot inflate its own yardstick, which is the failure of any score normalised by a locally estimated variance: such a score undershoots exactly where it should overshoot. This is what makes an unsupervised geometric test possible. A directional variant resolves that distance into a component along the animal’s local travel axis, *η_‖_*, and one across it, *η_⊥_*, which separates outliers that overshoot a real movement from outliers that displace the animal sideways, and is what the error classes are built on (Supplement S1).

### The probability detector

scores each location by how likely its speed, its turning rate and their step-to-step changes are under the animal’s own movement, so what counts as anomalous is set by the track rather than by a species constant. It divides those step-to-step changes by a scale that grows with the time gap, so irregular sampling is absorbed rather than penalised.

### The speed cap

does two jobs, which the cascade keeps apart, and two terms are worth separating before they are described: the *speed cap* is the detector, and the *speed ceiling* is the maximum step speed it may be given. Inside the combination the cap is state-relative and needs no ceiling: a step is flagged when its implied speed is anomalous against the local distribution, so 14 m s*^−^*^1^ is suspect within a rest segment even for an animal capable of 36. Given a ceiling, through mass and mode (Hirt *et al*., 2017) or found from the data, the cap instead fixes which pairs of locations a real step could connect, which is what the segmentation step needs.

### The detour detector

compares the path length between two locations with the straight-line displacement, so it needs no estimate of how far the animal should have moved. It is for the same reason insensitive to time: at sparse sampling it cannot separate a legitimate fast excursion from an erroneous one, and its threshold is the one constant in the cascade fixed in advance rather than read off the track.

### Thresholds read off the track

Key to unsupervised filtering is the fact that every threshold comes from the data, not the user. Existing tools ask for a maximum plausible speed, a distance limit, a turn-angle cut-off or a quality score, each depending on species, sampling regime, tag hardware and study question, and each is often guessed. Here, each detector instead scores every location and asks whether those scores separate into a bulk of ’ordinary movement’ and a tail of suspect points, placing its cut in the gap where one exists. The constants that govern the search are package defaults, exposed as arguments but not set by mt_clean_track() itself; Supplement S3 catalogues each with its classification and plausible range.

Alone, the detectors need not be silent, not even on clean data. Run on the clean reference track of Section 3.1 at the settings the cascade gives them, the probability detector flagged 19 of 3,537 locations and the detour detector 5, while the bridge detector and the speed cap flagged none; the detour detector’s own standalone default is more sensitive still, at 13. Over one hundred replicate realisations of that track, each carrying 30 m of per-location positional error, the cascade, however, raised only one false positive in total (Supplement S7.2). Because its consensus-finding rule refuses uncorroborated evidence, despite the detectors firing, it flags only exceptionally of the clean track’s locations.

### Combination by corroboration

The rule that turns four verdicts into one decision requires agreement rather than a majority. A majority vote, which could be a valid alternative consensus under certain conditions, would treat the detectors as interchange- able witnesses, which they are not: the detectors examine different properties of the same location, and one falling silent says nothing about the other. The default rule, evidence_corroborated, adds up each detector’s calibrated, signed evidence and flags a location when the total is positive and at least two detectors contribute to it. A single extreme detour witness may also act alone. The same evidence labels every flagged location with the kind of error it represents, the error_class fingerprint of Figure 1d, which the alternative class_aware rule uses as the decision itself.

### Block expansion

A spoofed or reflected signal produces a block of wrong locations, a run of them close together at a position the animal never occupied. Every step within that block is short and consistent with ordinary movement, so the detectors find nothing inside it. What they do catch are its seams, the two joins where the block meets the real track: the step out to the false position and the step back. In instances where the block lies mid-track those two seams suffice, because the real track continues on either side of it and each side establishes where the animal was. Successive passes of the cascade then work inward from both seams flagging every location with every pass. Where the block lies at the start or the end of a track however there is only one seam, with real track on one side of it and the block on the other, both internally ordinary. Nothing in the trajectory distinguishes them, and they remain invisible to the detectors. What I use here instead is an assumption, which this step relies and acts on: that most of a track’s locations are good quality real locations. After each pass the cascade therefore cuts the track into segments wherever no physiologically possible step could connect two consecutive locations (Figure 1c), retaining the largest segment and flagging every location outside it. A flag placed on a seam thereby extends to the whole block behind it, which is what the step’s name describes. On the cohort of Section 3.1 it flagged nothing, because CPF_D’s block lies mid-track and the detectors had already caught all of it before. Supplement S4.2 exercises it on a boundary spoof and Supplement S5.5 sweeps its scope. Outliers at the start and end of a track therefore remain the harder case, even where this step applies.

## 3 Evaluation

### 3.1 The cohort and the tools compared

I evaluated the cascade on a cohort of six synthetic tracks, bundled with the pack- age as inst/extdata/synthetic_tracks.csv.gz and reproducible end to end from inst/extdata/simulate_synthetic_data.R. Synthetic data are used because the evaluation needs to know which locations are outliers, which no real deployment could have supplied. Each track is drawn from a continuous-time movement model (Calabrese *et al*., 2016) and then contaminated with outliers of a known kind at known positions. The six carry the error morphologies the method is designed for, one dominant morphology each, and CPF_A to CPF_D sample irregularly at a nominal 15-minute interval, with missed locations and overnight gaps:

CPF_A 1,748 locations, 23 injected. Isolated spikes across four severity classes, plus a short burst.
CPF_B 3,537 locations, none injected. The clean reference, which measures false positives.
CPF_C 185 locations, 4 injected. A short, sparsely sampled track.
CPF_D 1,699 locations, 30 injected. A sustained spoof block of 30 contiguous locations, offset 800 km from the legitimate trajectory.
CPF_E 1,440 locations, 80 injected. Single-location radial excursions around a colony, at regular hourly sampling.
CPF_F 2,880 locations, 4 injected, at a regular 15-minute interval. A two-state movement process whose anomalies are kinematically ordinary outside their own state.

The CPF prefix marks the central-place forager process the first three tracks share and is retained as a label for the other three, which use different processes. Figure 2 maps each track with the flagged locations overlaid.

**Figure 2:**
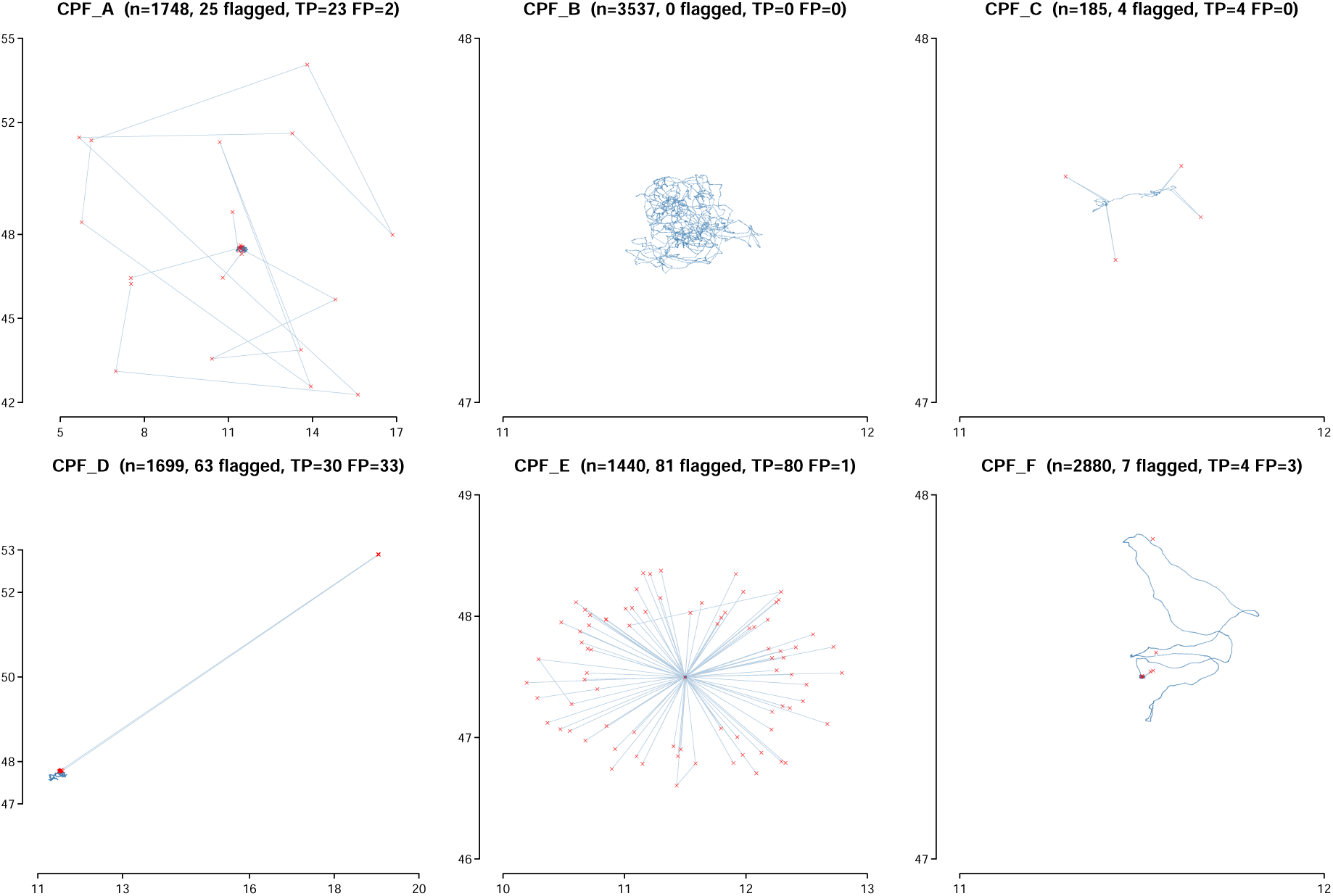
Cleaning-cascade outcomes across the synthetic cohort (CPF_A– F; the six error morphologies are described in **Section 3.1**). Each panel maps one track’s raw trajectory in steel-blue with the locations the zero-parameter cascade flags overlaid as red crosses; axes are longitude and latitude in degrees (WGS84).Panel titles give the track ID, location count (*n*), total locations flagged, and true and false positives (TP, FP) against the bundled ground-truth outlier set. Generated by scripts/worked_examples.R.

The synthetic data serves mainly to compare the performance of the algorithms against each other. What is more, the performance is specific to the movement process chosen and would not necessarily translate to performance in relation to real data. Real data, as I mentioned before, suffers from the fact that there is no way to know what constitutes a real outlier. Comparing between algorithms on such data would hence be confounded by the many dimensions that affect measured performance, all of them weighed in ignorance of what constituted an outlier in the first place.

I ran four competitors against the same cohort. A naïve fixed speed cap, a one-line dplyr::filter(speed < v_max), gave the floor case. atlastools (Gupte *et al*., 2022) flags a location only when both its incoming and its outgoing speed exceed the threshold, and is pinned by commit (pratikunterwegs/atlastools@3ba76c5) since it is not on CRAN. The package also ships a dedicated filter for prolonged spikes, which cannot be swept over a dataset unsupervised and is reported separately in Supplement S4.3. trip::sda (Freitas *et al*., 2008) is the Freitas speed, distance and angle filter, and SDLfilter::ddfilter (Shimada *et al*., 2012) adds angle-aware loop detection to a speed cap; both ran at their documented defaults. The comparison I chose is deliberately asymmetric in the competitors’ favour. Each of them requires a maximum plausible speed, and each was given 1.5 m s*^−^*^1^, twice the fastest step in the clean reference track: a threshold read off data known to contain no outliers, which no real deployment provides. The cascade in contrast was given the track and nothing else, as mt_clean_track(x). Two measures score every tool below and answer different questions: recall is the share of a track’s (injected and thus known) outliers that a tool flags, precision the share of its flags that were actual outliers. *F*_1_, the harmonic mean of those two, summarises them in one number that is high only when both are high; Supplement S4.1 defines all three.

The function mt_clean_track() can also be given a behavioural state vector or a physiological speed bound. Either argument should only be supplied when the track has the problem it is intended to solve. Only CPF_F carries either, and neither improved this cohort: the state vector left CPF_F unchanged in Table 2 and cost precision over replicates (Supplements S5 and S7.2).

**Table 2:** Detection skill per track for the cascade and four competitors. Each cell is the per-track *F*_1_ score against the track’s bundled truth set (Supplement S4.1): 1.00 is perfect recovery with no false positives, and higher is better. CPF_B carries no injected outliers, so *F*_1_ is undefined and that row reports each tool’s false-positive count instead (0 is ideal). The competitors, and the single ceiling *v*_max_ every tool was given, are defined in Section 3.1. **cascade** is mt_clean_track() at its zero-parameter default; **cascade (+state)** the same call with a behavioural-state column supplied, which only CPF_F carries, so elsewhere that cell reads “—”. Every row comes from one run of scripts/cohort_competitors.R against move2utils v0.4.5.

| Track | naive | atlastools | trip::sda | SDLfilter | cascade | cascade (+state) |
| --- | --- | --- | --- | --- | --- | --- |
| CPF_A | 0.852 | 0.978 | 0.955 | 0.979 | 0.958 | — |
| CPF_B (clean) | 0 FP | 0 FP | 0 FP | 0 FP | 0 FP | — |
| CPF_C | 0.667 | 1.000 | 1.000 | 1.000 | 1.000 | — |
| CPF_D (spoof block) | 0.062 | 0.000 | 0.065 | 0.176 | 0.645 | — |
| CPF_E (colony halo) | 0.675 | 0.976 | 0.994 | 0.976 | 0.994 | — |
| CPF_F (multi-state) | 0.500 | 0.667 | 0.667 | 0.889 | 0.727 | 0.727 |
| Mean $F_1$ (truth tracks) | 0.551 | 0.724 | 0.736 | 0.804 | **0.865** | 0.727 |

### 3.2 The coherent block

On the sustained spoof block of CPF_D all four competitors failed but the cascade did not: it recovered all 30 injected locations, where SDLfilter recovered three, trip::sda and the naïve cap one each, and atlastools none. The recovery was not the work of the block-expansion step, which declined because the block lies mid-track (Supplement S7). It was the work of the iteration loop, which peeled the block inward from its seams over 28 passes, recovering a single injected location on 27 of them and three on one, accounting for all 30. Each individual flag is decided by corroboration among the detectors, and the block as a whole is removed from the track by repetition. Sweeping the ceiling from 0.5 to 36 m s*^−^*^1^ did not improve the performance of the competitors as none reached the parameter-free cascade at any value (Supplement S5).

Since atlastools requires both the incoming and the outgoing speed of a location to be extreme, and no interior location of a coherent block satisfies that, it hardly recovered any of the outliers. The package also ships a dedicated filter for prolonged spikes, which recovered all 30 locations, but it has to be aimed at a block which somehow has already been seen, and on the tracks that carry none it failed outright (Supplement S4.3).

That result is specific to the block this cohort injects. A 30-location block was recovered wherever it lay. A longer one was not. While a 300-location block was recovered in full at or near either end of the track, in the middle it was recovered only partly, in the worst case 13 of 300 locations (Supplement S5.5).

Both steps in the cascade that catch a block run out on a long one in the middle of a track. Block expansion cuts the track wherever no animal could have travelled and keeps only the largest piece. Near a track’s end the rest of the track is still one large piece, so the step acts. In the middle the same block leaves two halves of similar size. Neither is obviously the real track, so the step holds back rather than discarding genuine movement. That leaves the iteration loop working alone. It strips a block from its seams and stops after a hundred passes. Thirty locations take twenty-eight passes and finish. Three hundred do not.

Two things follow for anyone expecting long runs of false positions. First, the cleaned track records why the cascade stopped: when it stopped at the pass limit rather than for want of anything left to flag, it says so, and a half-removed block can be told from a clean one. Raising that limit lets it carry on. Second, a block that grows to rival the real track in number of locations cannot be removed by any setting.

### 3.3 Across the cohort

The cascade left no injected outlier behind on any track. Of the 141 outliers injected across the five contaminated tracks it recovered all, at a recall of 1.000 on every one, while the competitors left between 27 and 33 behind. Every point on which it scored below a competitor was a cost in precision, not a missed outlier (Table 2).

*F*_1_, even though I chose this as a performance metric, is not truly an honest summary. It weights a false negative and a false positive equally, and in an analysis pipeline, downstream they are not equal. A retained outlier has the potential to displace whatever is estimated from the track: one location 800 km away moves a home-range polygon, inflates a step-length distribution and creates a speed no animal produced. A discarded true location, a false positive, costs a little data, and on these test tracks the cascade discarded a few dozen from thousands. The asymmetry follows from the assumption on which the whole approach rests, which is that outliers are a minority of the track. That assumption is what lets a threshold be read off a score distribution at all, and the same assumption makes a false positive cheap: it is drawn from the large majority of ordinary locations. A false negative is drawn from the small minority, and its effect on whatever is estimated next is proportionally much larger. The two tracks where the cascade scored below SDLfilter were both of this kind: it matched that tool on recall yet paid in precision.

Averaged over the single realisation of Table 2, the cascade led on mean *F*_1_ at 0.86 against 0.80 for the strongest of the four, and it was the one tool that was not told anything. Pooling the counts rather than averaging the tracks narrowed that to 0.88 against 0.87, because pooling weights each track by the outliers injected into it.

Over one hundred replicate realisations, each carrying 30 m of per-location positional error, a competitor took the higher mean *F*_1_ on three of the five contaminated tracks, in each case with non-overlapping intervals, and each worked from a ceiling calibrated on the clean reference track that the cascade never received. The cascade’s recall never fell below 0.958, so what it gave up was precision, not outliers. What the competitors gave up was outliers: all four collapsed on the coherent block, to 0.160 recall or below, and the naïve cap led nowhere. Across the cohort the cascade still led on mean *F*_1_, 0.821 against 0.777 (Supplements S7.2 and S5.4). CPF_F is the only track carrying behavioural states, and supplying them did not help it. Over the same one hundred replicate realisations the cascade scored a mean *F*_1_ of 0.954 with the states supplied against 0.956 without them, and with 30 m of positional error 0.468 against 0.616, at recall unchanged in both settings: what the states added was false positives rather than recovered outliers.

Choosing the competitor that suits a track requires knowing which error morphology it carries, which is what a screening step is there to establish, and where a deployment carries several the choice cannot be made at all.

### 3.4 Caveats

The cohort is built from one stochastic family, Ornstein–Uhlenbeck foragers from ctmm::simulate() with controlled outlier injection. It has no behavioural regimes richer than CPF_F’s two-state concatenation, no multiple error sources co-present in one deployment (atmospheric drift, multipath, hardware drift and spoofing together), and no sampling messier than the simulated duty cycling. The cohort therefore tests the floor case for the cascade.

Every number should be read as indicative rather than definitive in the evaluation. The comparison is also deliberately tilted against the method being proposed. The competitors were handed a speed ceiling calibrated on a track known to contain no outliers, the best case their tuning admits and it is something that is not available in real use-cases, while the cascade was given the track alone. Nothing in the cascade was tuned on this cohort: its constants are package defaults, fixed before these tracks were generated. The figures therefore support a claim about ordering under conditions favourable to the alternatives, not about the magnitudes themselves.

Detection outcome here means the four cells of a confusion matrix: true positives, false positives, false negatives, and the true negatives that go unreported because they dominate every track by construction. Outliers are rare by definition, and that rarity is why precision and recall rather than accuracy are the axes used throughout.

That same rarity is where the approach breaks down. A screen that infers its thresholds from a track’s own score distribution needs the anomalies to be a minority of that distribution. When they are not, the contamination defines the bulk, and the cascade stops early on a track it has barely cleaned. This is a general property of unsupervised outlier detection.

## 4 Adapting to real data

The default, a single-track move2 object with no further arguments, covers most use, though the function would also handle multi-track move2 objects. One step belongs before the cascade, and four situations call for more from it, each easier to recognise by its symptom than by the argument that answers it.

### Preflight preparation

Some locations should never reach the cascade, and none of them is a trajectory problem. mt_filter_gps_quality() drops locations on the fields a tag records about its own fix rather than on the path they describe: satellite count, dilution of precision, horizontal accuracy, and geometries that are empty because no location was obtained at all. It thereby reaches the one class of error no trajectory-based test can, cold-start acquisition, which is indistinguishable from fast genuine movement. Deployment records answer a second class. A location at the very start or the very end of a track, sitting far from everything else, is usually the tag reporting before it was attached or after it came off, and no per-location test can see it, there being no bracketing neighbourhood against which it is anomalous. Trimming a track to the interval over which the tag was on the animal removes such locations by knowing something the trajectory does not contain.

### A track too short to fit its own thresholds

A detector that reads its threshold from a score distribution needs enough scores for the cut to be visible. pool_by fits the thresholds on a group of tracks, an individual, a study or a species, and unions the flags this adds back into each track, so that a short deployment borrows the structure of the longer ones it belongs with. A two-level c(outer, inner) form is for when the unit the thresholds are fitted on and the unit they operate on differ. Pooling across a group (outer) is a first step towards a hierarchy in which an individual (inner) borrows what it lacks from the deployments it came with, since what is ordinary for an animal can be assumed to be in part what is ordinary for animals like it.

### Behaviour that changes within a track

A track mixing rest and flight carries two step-length distributions, and a threshold fitted on their union sits between them: too loose for the resting stretch and too tight for the flying one. state takes the state the animal was in at each location, from a hidden Markov model or assigned by an expert, and runs the kinematic detectors within each behavioural segment instead. A global rescue pass keeps the geometric detectors on the full track at the same time, so a per-state threshold cannot dilute geometric evidence. Whether this improves anything depends on the deployment: it did not on the cohort here, where the over-flagging turned out to be geometric (Section S1.9).

### Positions no animal could have reached

Supplied through the animal’s mass and locomotion mode (Hirt *et al*., 2017), a physiological speed bound gives the cascade an absolute ceiling on how fast a real step can be, and so tells it which pairs of positions a real animal could have connected. That makes two structural steps possible: peeling off locations that no real step could have reached, and cutting the track into reachable segments, so that an unreachable block is identified as a whole rather than location by location.

Smaller options supply an observation-error prior (location_error), asymmetric pre- peeling and class-aware persistence filtering. The mechanics and worked examples are in Supplement S1.11 and the package vignettes; the whole procedure is set out as pseudocode in Algorithm 1 (Supplement S2), and every constant is catalogued in Supplement S3.

## 5 Discussion

Animal paths are read for what they say about physiology, behaviour and the use of landscapes, and every one of those readings inherits the positions it starts from. Screening those positions is therefore not a preliminary to the analysis but part of it. A screen that reads its reference off the track removes the analyst’s arbitrary choice without removing the evidence: each flag carries the kind of error it represents and the evidence that produced it, so the decision stays open to inspection rather than buried in a filter. The gain is not only cleaner tracks but tracks whose cleaning is itself part of the record.

Every method that removes outliers from a movement track must first answer the question: against what reference is a location judged? The existing answers are usually borrowed from outside: a speed filter takes its reference from the species and the device, a state-space or continuous-time model from an assumed movement process, and a calibration-based diagnostic from a test deployment whose error is known, or the number of satellites used to derive location. Each is a sound statement about an animal, a tag or a process, and usually nothing in the procedure reports when they have diverged from the track in hand. What I propose is to take the reference from the track itself. External knowledge is not banished, since a physiological ceiling enters wherever mass and locomotion mode can be supplied (Section 4), or an initial cleaning based on number of satellites clearly improves the cleaning process. But what I argue for is that this knowledge enters as a declared bound rather than an unexamined default, a fact that can be tested.

That bears on an understated distinction the field draws constantly. Every location carries error, because that is what a measurement is; an outlier is not a location with error but one whose error is large enough that keeping it would distort insight. The boundary between them is a decision and not a discovery, and the whole difficulty sits in deciding which are too far out to keep. Reading the reference off the track does not abolish that convention, but it moves the convention out of the analyst’s prior and into the animal’s own record. What two deployments then share is not a threshold but the rule that produced one, since a single number is the wrong quantity to carry across studies.

Whatever a method measures, and however well founded the measurement, it usually ends by handing the analyst a number and a question: should I keep this location, or drop it? A speed cap answers with a limit fixed in advance; a likelihood or a residual under a fitted movement process answers far more finely, grading each location by how improbable it is, often leaving the removal to the analyst’s judgement. The measurement has, I would argue, become principled while the cut has not, leaving some residual arbitrariness. What I propose is that the cut come from where the scores come from: the bulk of a track’s scores describes what that animal ordinarily did, and the point at which that bulk ends is a feature of the distribution rather than a preference of whoever reads it, so it can be located rather than chosen.

What locating a cut in a measurement leaves open is worth separating from what it closes. Where a distribution’s bulk ends is a property of the data; how much corroboration to require before dropping a location is not. A threshold filter cannot hold the two apart, its threshold being at once the evidence and the decision. Here a detector supplies evidence on a scale it shares with the others and a combination rule turns that evidence into a decision, so either can change without redesigning the other. The four detectors are ingredients to the recipe of the default rule, and not the method itself. What the cascade returns is therefore the foundational evidence, which makes it a screening step and not an error model.

An internal reference is bought at a price. A track can supply a reference only if it contains enough of its own normality to define one, which is the condition Section 3.4 sets out and the point at which the approach fails. Where the track cannot supply a solid reference, it has to be provided: behavioural states supply the structure a mixed track lacks, pooling in cases where a short deployment has too little data, and a physiological ceiling a bound no quantity of data contains.

### 5.1 Outlook

Because the cascade returns per-location flags and a continuous combined_evidence score, its output could in theory serve an analysis that prefers to retain and down-weight noisy locations rather than delete them, as an observation-quality input for example to a continuous-time smoother (Johnson *et al*., 2008; Calabrese *et al*., 2016; Jonsen *et al*., 2023). The clearest room for improvement is the combination rule, which weights the detectors equally, a loss if they carry asymmetric information. Exposing the combiner as an interface taking explicit weights or an arbitrary function would turn the tolerance above into a setting rather than a consequence, and rules learned from labelled tracks, including neural approaches to consensus among detectors, could adapt that weighting to the deployment.

Four detectors and one combination rule are not the end of this. What I have set out is a way of asking the question, in which the track supplies the reference, the data locates the cut, and the evidence outlives the decision it informs, so that better detectors and better rules can be substituted without rebuilding the argument around them. As tags multiply and new sources of error appear, from hardware drift to deliberate interference with the satellite signal, so will more sophisticated detection algorithms; transparency and reproducibility will matter more than any particular set of thresholds.

## Data availability

This paper uses no observational data. Every track analysed is synthetic and is regenerated deterministically, seeds included, by move2utils/inst/extdata/ simulate_synthetic_data.R, which uses the ctmm package (Calabrese *et al*., 2016) for the OUF generative model and records all model parameters and ground-truth outlier positions; the copies shipped as move2utils/inst/extdata/synthetic_ tracks.csv.gz and move2utils/inst/extdata/synthetic_ground_truth.rds are a convenience, not the source of record. The cohort map of Section 3, and the per-track metrics and flag-class composition of Supplement S7, are emitted by scripts/worked_examples.R; the multi-tool comparison tables (Section 3.2, Supplement S4.4) by scripts/cohort_competitors.R, which computes every row, competitors and cascade alike, against move2utils v0.4.5 in a single run; the ceiling, sampling and positional-error experiments of Supplement S5 by scripts/vmax_sensitivity.R, scripts/sampling_sensitivity.R and scripts/error_sensitivity.R; the worst-case summary of Supplement S5.4 by scripts/robustness_summary.R; the replicate intervals of Supplement S7.2 by scripts/replicate_ci.R, with the cohort-shape comparison quoted there by scripts/cohort_shapes.R; the wall-time scaling figure (Supplement S6) by scripts/runtime_scaling.R; the synthetic block-expansion demonstration (Supplement S4) by scripts/block_demo.R. The move2utils package is openly available at https://github.com/move2universe/move2utils. The version this paper describes is 0.4.5, as recorded in the package DESCRIPTION; the archived deposit below is the citable record for it. The analysis scripts, the synthetic-track generator and the emitted results are archived on Zenodo at https://doi.org/10.5281/zenodo.22080020. That deposit is the citable record from which every figure and table in this paper can be reproduced.

## Conflict of interest

The author declares no competing interests.

## Funding

No specific funding was received for this work.

## Use of generative AI

I used Claude Code (Anthropic; Claude Opus 4- and 5-series models, 2026) during the preparation of this manuscript, where the tool assisted with drafting, editing, and grammatical correctness, and with cross-checking claims in the text against the move2utils package source code. Literature search and verification of all cited references were performed by me without AI assistance. The same tool was used during development of the move2utils package by its authors (Kranstauber, Safi, Scharf), where it assisted with drafting and refactoring R code, generating scaffolding for unit tests, and surfacing legacy inconsistencies across exported functions; AI-assisted code was reviewed, tested, and integrated by the package authors and is traceable through the package’s git history. I reviewed all AI-generated material in this manuscript, take full responsibility for its content, and confirm that no AI tool is listed as an author.

## S1 Methods in detail

### Notation

Symbols recur across the sections below, and a reader entering the supplement at any one of them will need this table rather than the section that first defines them. Each symbol is defined once, here and where it is introduced, and is used in the same sense throughout.

**Table 3:** Symbols used throughout the supplement.

| Symbol | Name | Meaning |
| --- | --- | --- |
| $\mathbf{x}_i, t_i$ | position, time | the $i$th location of a track of $n$ locations, and its timestamp |
| $\tau_i$ | time lag | $t_{i+1} - t_i$, the interval to the next location |
| $d(\cdot, \cdot)$ | distance | geodesic distance between two positions |
| $v_i$ | step speed | $d(\mathbf{x}_i, \mathbf{x}_{i+1})/\tau_i$ (Eq. 1) |
| $\theta_i$ | turning angle | angle between successive displacement vectors |
| $\omega_i$ | angular velocity | $\theta_i/\tau_i$ (Eq. 2) |
| $\Delta v_i, \Delta \omega_i$ | auto-differences | change in speed and in angular velocity between successive steps (Eq. 3) |
| $P_{\text{st}}$ | step-turn probability | joint density of $(v, \omega)$, estimated from the track itself |
| $P_{\text{joint}}$ | joint probability | the three components combined (Eq. 4); the probability detector's score |
| $p_i$ | probability evidence | signed evidence contributed by the probability detector |
| $r_v, r_\omega$ | lag-1 autocorrelations | persistence of speed and of angular velocity |
| $\alpha$ | persistence weight | how far the auto-difference components are trusted, from $r_v$ and $r_\omega$ (Eq. 5) |
| $s(\tau^\Delta)$ | gap-dependent scale | scale by which an auto-difference is normalised for the gap it spans |
| $\mu_i$ | bridge mean | expected position of location $i$ given its neighbours and the elapsed times (Eq. 10) |
| $\sigma_i$ | bridge width | the spread expected at that point, built from the timestamps alone (Eq. 11) |
| $\eta_i$ | bridge residual | $\ \mathbf{x}_i - \mu_i\ /\sigma_i$ (Eq. 12); the bridge detector's score |
| $\eta_{\parallel}, \eta_{\perp}$ | directional residuals | the residual resolved along and across the local travel axis (Eq. 13) |
| $\rho_k$ | detour ratio | path length divided by displacement over a window of $k$ steps (Eq. 15); the detour detector's score |
| $c$ | speed cap | the step speed above which a location is flagged (Eq. 16) |
| $c_{\text{phys}}, v_{\text{max}}$ | speed ceiling | an absolute physiological bound, supplied through `mass` and `mode` or estimated allometrically (Eq. 17) |
| $T$ | track | the input track, of $n$ locations |
| $k$ | window or component size | the number of steps in a detour window, or of locations in a connectivity component |
| TP, FP, FN | flag counts | an injected outlier flagged, a legitimate location flagged, and an injected outlier left in place (Eq. 21) |
| $F_1$ | detection score | the harmonic mean of precision and recall (Eq. 22) |

### S1.1 The probability detector

The core idea is to ask, for each location in a track, how probable the observed movement is under the empirical distribution of the track as a whole. An animal moving normally produces a characteristic signature in the joint space of speed, angular velocity, and the persistence of these quantities between consecutive steps. An outlier disrupts this signature: the speed to and from it is anomalously high, the angular velocity implies an abrupt directional reversal, and the changes in speed and turning rate between consecutive steps are discontinuous. By quantifying how probable each location’s movement signature is, the method identifies locations that are inconsistent with the animal’s own behaviour without requiring any prior specification of what “normal” looks like.

Consider a track of *n* locations **x**_1_*, . . . ,* **x***_n_* recorded at times *t*_1_*, . . . , t_n_*. For each location *i* (2 *≤ i ≤ n −* 1) I compute three probability components. The first component is the **step–turn probability** *P*_st_(*v_i_, ω_i_*), which captures how likely the observed combination of speed *v_i_* and angular velocity *ω_i_* is under the joint empirical distribution. Speed is defined as

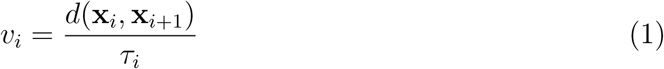

where *d*(*·, ·*) is the geodesic distance and *τ_i_* = *t_i_*_+1_ *− t_i_* is the time lag. Angular velocity is

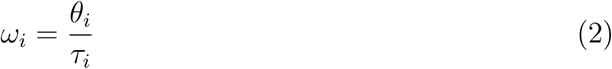

where *θ_i_* is the turning angle between the displacement vectors **x***_i_ −* **x***_i−_*_1_ and **x***_i_*_+1_ *−* **x***_i_*. Dividing by *τ_i_* is what makes the score comparable across gaps: a displacement of 500 m in one hour and 500 m in one minute represent fundamentally different movement states, yet classical step-based approaches treat them identically. By operating on speed and angular velocity rather than raw step lengths and turning angles, the method is time-aware: irregular sampling is absorbed by the normalisation rather than penalised. Speed and angular velocity estimated from discrete locations are scale-dependent: with increasing time lags, the straight-line distance between locations increasingly underestimates the true path length, and derived speeds underestimate the true velocities (Rowcliffe *et al*., 2012; Noonan *et al*., 2019). For regularly sampled data, this normalisation rescales uniformly and does not affect relative probabilities.

The joint density *P*_st_(*v, ω*) is estimated as a two-dimensional histogram using the Freedman–Diaconis rule for bin widths (Freedman & Diaconis, 1981). Because the angular axis is circular, the histogram is padded by tiling copies at *±*2*π* before bilinear interpolation smooths the surface. Each location’s probability is then extracted from this smoothed 2D density. Alternatively, a parametric variant fits Weibull marginals to speeds (Morales *et al*., 2004) and von Mises marginals to angular velocities (Mardia & Jupp, 1999) and returns the product of the marginal densities. The parametric approach is faster and can perform better with small samples, although it assumes unimodal marginals.

The second and third components capture the **persistence** of movement, typical for all animal movement (Gurarie *et al*., 2017), and represent how much speed and angular velocity change between consecutive steps. An animal moving steadily produces small changes between successive measurements; an outlier inserted into the track produces a spike in speed followed by an equally abrupt deceleration, creating large consecutive changes that a real animal would rarely exhibit.

I define the auto-differences

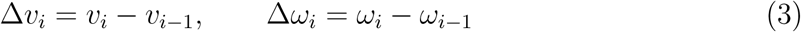

and estimate their marginal densities *P* (Δ*v*) and *P* (Δ*ω*) via kernel density estimation (Silverman, 1986). The delta-speed component captures whether the acceleration or deceleration at a given location is unusual relative to the track’s overall dynamics. The delta-angular-velocity component captures unusual shifts in turning rate: for instance, the abrupt directional reversals often produced by GPS measurement error (Hurford, 2009).

The joint probability at location *i* combines all three components:

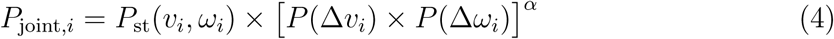

The exponent *α* controls the relative weight of the auto-difference terms. Its value determines how much the persistence components contribute to the joint probability: *α* = 0 ignores auto-differences entirely (relying on speed and angular velocity alone), while *α* = 1 gives them equal weight with the step–turn component.

How informative the auto-differences are depends on the autocorrelation structure of the movement. When speed is highly autocorrelated, the animal maintains its pace over consecutive steps, a sudden change in speed is a strong signal of an anomaly, and the auto- difference should carry weight. When speed has low autocorrelation (because the animal switches behavioural states frequently, or because the sampling interval is long relative to the decorrelation time), consecutive speed measurements are nearly independent, and the delta-speed distribution is broad even for normal movement. In this case, the auto- difference adds noise rather than signal. Angular velocity exhibits the same pattern, often with substantially lower autocorrelation than speed.

I set *α* from the data by computing the lag-1 autocorrelation of speed (*r_v_*) and angular velocity (*r_ω_*) and taking the geometric mean:

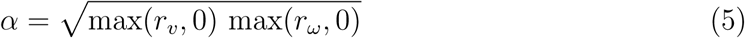

This ensures that *α* is high when both components carry persistent, informative signals, and low when either is dominated by noise. On the synthetic cohort the estimator returns *α* between 0 and 0.34, so the auto-difference term carries modest weight throughout. On five of the six tracks the reason is the one the estimator exists for: speed is persistent between consecutive steps while angular velocity is close to independent, and the estimator declines to let an uninformative angular channel dilute the joint probability. On CPF_D it is the other way round, and the consequence is stronger than downweighting, and it is a defect of the estimator rather than a property of the track. There the lag-1 autocorrelation of speed is itself zero, plausibly because the injected 800 km excursions destroy step-to-step persistence: so *α* = 0 exactly, the bracketed term becomes identically 1, and the persistence components contribute nothing at all. The block is recovered there by the geometric detectors and the consensus-finding rule, and the persistence machinery, including the gap-aware normalisation below, is inactive while it happens. Evaluating that normalisation rather than describing it would need a track on which *α* is substantial, which this cohort does not provide. The estimator is also vulnerable to the contamination it exists to detect. Recomputing *α* on each track with the injected outliers removed raises it on every track where the comparison is possible (CPF_A 0.139 *→* 0.264, CPF_C 0.053 *→* 0.381, CPF_D 0.000 *→* 0.343, CPF_F 0.238 *→* 0.319), so the outliers reduce the weight given to the term meant to catch them. That is the masking effect the bridge’s timestamp-only denominator was designed to avoid, reappearing in a data-driven tuning parameter; computing *α* on a trimmed track, or after a first cleaning pass, would remove it.

The product formulation in Eq. 4 assumes independence among the three components. Although speed, angular velocity, and their consecutive changes are not strictly independent in animal movement, the product provides a useful composite score that, in practice, effectively separates outliers from the bulk of the data (Hand & Yu, 2001).

### S1.2 Gap-aware auto-differences

The time normalisation in Eqs. 1 and 2 makes the step–turn component robust to irregular sampling. The auto-difference components (Eq. 3), however, require a further correction. The change in speed Δ*v_i_* = *v_i_ − v_i−_*_1_ depends not only on the animal’s behaviour but also on the time gap *τ* ^Δ^ between the two steps being compared. For any movement process with velocity autocorrelation *ρ*(*τ* ), the expected squared auto-difference is

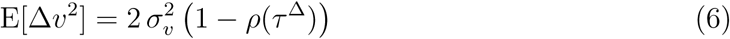

where *σ*^2^ is the stationary variance of speed and *τ* ^Δ^ = (*τ_i−_*_1_ + *τ_i_*)*/*2 is the time between the midpoints of the two steps (Cressie, 1993; Fleming *et al*., 2014). At short lags, *ρ ≈* 1 and speed changes are small: the animal’s velocity is autocorrelated and consecutive measurements are similar. At lags much longer than the decorrelation time, *ρ →* 0 and Δ*v* approaches the difference of two independent draws from the stationary distribution. With regular sampling this is immaterial: all gaps are equal and the scaling is uniform. With irregular sampling, the auto-difference distribution conflates measurements at different temporal scales. A large speed change after an overnight gap is normal (the animal’s velocity has decorrelated), yet it would be penalised by a density built predominantly from short-gap deltas. Conversely, a modest speed change after a sho brt gap may be genuinely anomalous but is diluted by the long-gap portion of the distribution.

I address this by estimating the gap-dependent scale of each auto-difference non- parametrically. The observed pairs (*|*Δ*v_i_|, τ* ^Δ^) are binned into equal-count groups by *τ* ^Δ^, and the median absolute deviation (MAD) is computed within each bin (Leys *et al*., 2013). Interpolation yields a smooth scale function *s*(*τ* ^Δ^) mapping any gap to the expected magnitude of the auto-difference at that gap. Each auto-difference is then normalised:

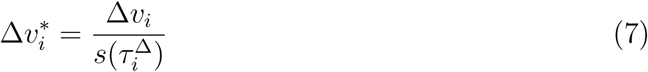

The kernel density estimate is built on the normalised values Δ*v^∗^*, and scoring applies the inverse transformation to recover densities in the original space:

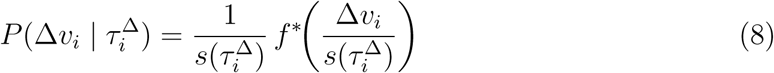

where *f^∗^* is the KDE of the normalised auto-differences. The same procedure applies independently to Δ*ω*. This approach makes no assumption about the functional form of *ρ*(*τ* ): for an Ornstein–Uhlenbeck process, *ρ* is exponential; for a migrating animal whose speed regime shifts over days, the autocorrelation structure may be entirely different. The non-parametric estimation adapts to whatever the data exhibit. For perfectly regular data, all gaps are equal, *s*(*τ* ^Δ^) is constant, and the method reduces identically to the original formulation.

The gap-dependent scaling has implications beyond outlier detection. Any method that constructs empirical distributions of consecutive changes in movement metrics from tracking data (whether for trajectory simulation, behavioural change-point detection (Gu- rarie *et al*., 2009), or path reconstruction), faces the same conflation of temporal scales when applied to irregularly sampled data. The decomposition into a gap-independent shape (*f^∗^*) and a gap-dependent scale (*s*(*τ* ^Δ^)) resolves this generally, allowing such methods to operate at arbitrary and variable time intervals while preserving the autocorrelation structure.

### S1.3 Separating outliers from the bulk

#### Entropy−valley thresholding is a no−op on clean data

Having assigned each location a joint probability *P*_joint_*_,i_*, the task is to determine where the boundary between outliers and legitimate movement lies. This is fundamentally a separation problem: the distribution of log *P*_joint_ across all locations in a track typically exhibits a characteristic structure. The bulk of the data, locations where the animal moved normally, clusters in a relatively narrow, high-probability region. Outliers, by contrast, fall in the far lower tail: their speed, angular velocity, or persistence is so unusual that their joint probability is orders of magnitude below the bulk. Between the two lies a transition zone whose sharpness depends on the severity of the outliers and the variability of normal movement.

The key insight is that this separation is a property of the data, not a parameter to be specified. Two locations whose log *P*_joint_ values differ by four orders of magnitude are unlikely to belong to the same distribution. The challenge is to identify this boundary automatically. I provide two complementary approaches, each suited to different distributional shapes.

#### S1.3.1 Gap-based thresholding

If the sorted log *P*_joint_ values arose from a single continuous distribution, the gap sizes between consecutive sorted values would follow a predictable pattern determined by the harmonic series, as in MacArthur’s broken-stick model for species abundance (MacArthur, 1957). I compute the ratio of each observed gap to its expected size under this null model. Gaps in the lower tail that substantially exceed their expectation signal a discontinuity between outliers and the bulk.

This initial signal is refined through tail-decay inflection analysis. Starting from the most extreme sorted value, I sequentially remove candidate outliers and track the resulting change in tail length: the distance from the current leftmost value to the edge of the bulk distribution. Removing genuine outliers produces steep drops in tail length; upon entering the bulk, these drops become small and approximately linear. The inflection point, where the second derivative of the tail-length curve drops to the noise floor, marks the natural boundary.

This two-stage approach makes no distributional assumptions about the shape of the log *P*_joint_ distribution, requires no user-specified threshold in the statistical sense (only a sensitivity parameter governing the gap ratio, with a robust default), and converges naturally under iteration because removing outliers eliminates the break that triggered their detection.

#### S1.3.2 Density-valley thresholding

An alternative approach exploits the density structure of the log *P*_joint_ distribution directly. I estimate the density of all valid log *P*_joint_ values by Gaussian kernel density estimation, at R’s default rule-of-thumb bandwidth, and search for local minima (valleys) below the main mode. The bandwidth is the procedure’s operative parameter and is not tuned: whether a valley exists at all is decided by it, and because the rule-of-thumb bandwidth shrinks with *n*, the same contamination fraction can separate at one track length and not at another. A valley indicates a natural separation between two regimes: an outlier regime in the lower tail and the bulk distribution of normal movement (Silverman, 1981). The deepest valley whose density falls below a fraction *δ* of the peak density (default *δ* = 0.3, unified package-wide across the probability, bridge, and speed cap detectors; see Supplement S3) is taken as the threshold:

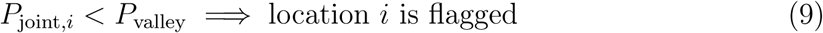

If no valley meets the depth criterion, the distribution is effectively unimodal and no outliers are declared. This approach is more sensitive than the gap method in practice, because it identifies the boundary between regimes from the shape of the entire density rather than from a single gap in the ordered sequence. It is particularly effective when outliers span a range of severities, producing a broad lower tail rather than a sharply separated cluster. Two further threshold methods are available for users who prefer conventional approaches: a significance method based on robust *z*-scores (median and MAD; Leys *et al*., 2013) of log-probabilities, and a fixed-percentile method.

### S1.4 The bridge detector

Figure 4 illustrates the construction: the bridge interpolated between a location’s neighbours, the residual it leaves, and the directional decomposition that separates displacement along the expected path from displacement across it.

**Figure 3:**
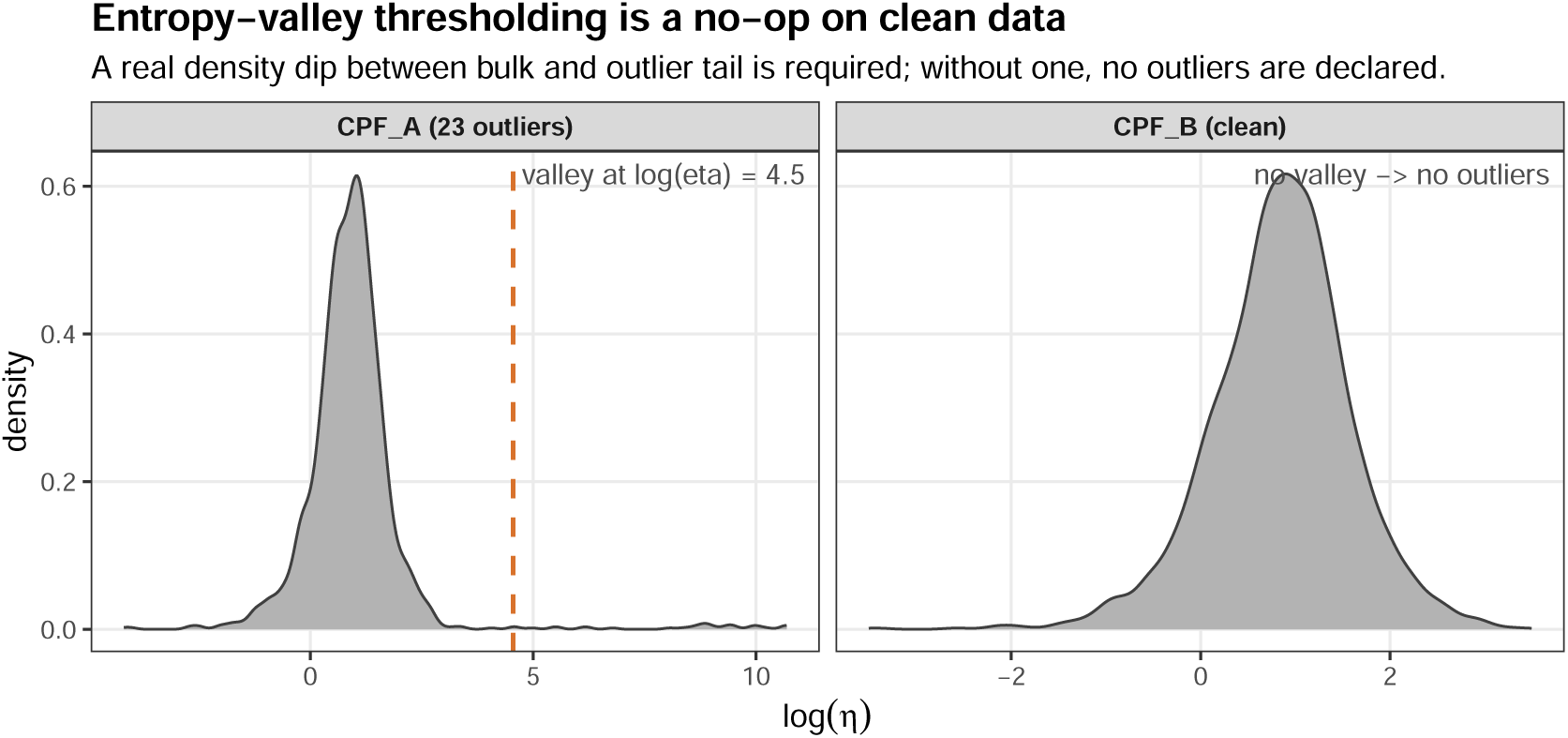
Entropy-valley thresholding is a no-op on clean, regularly sampled data. Left: density of log(*η*) for CPF_A (1,748 locations, 23 injected outliers) shows a clear bimodal shape with a valley separating the bulk of normal movement from the outlier tail. The density-valley detector places the threshold (dashed red line) in the valley. Right: density of log(*η*) for CPF_B (3,537 clean locations) is smoothly unimodal with no valley, and the detector correctly returns no outliers. A gap-detection threshold applied to the same distributions, by contrast, fires on the natural tail of CPF_B and produces false positives on clean data.

**Figure 4:**
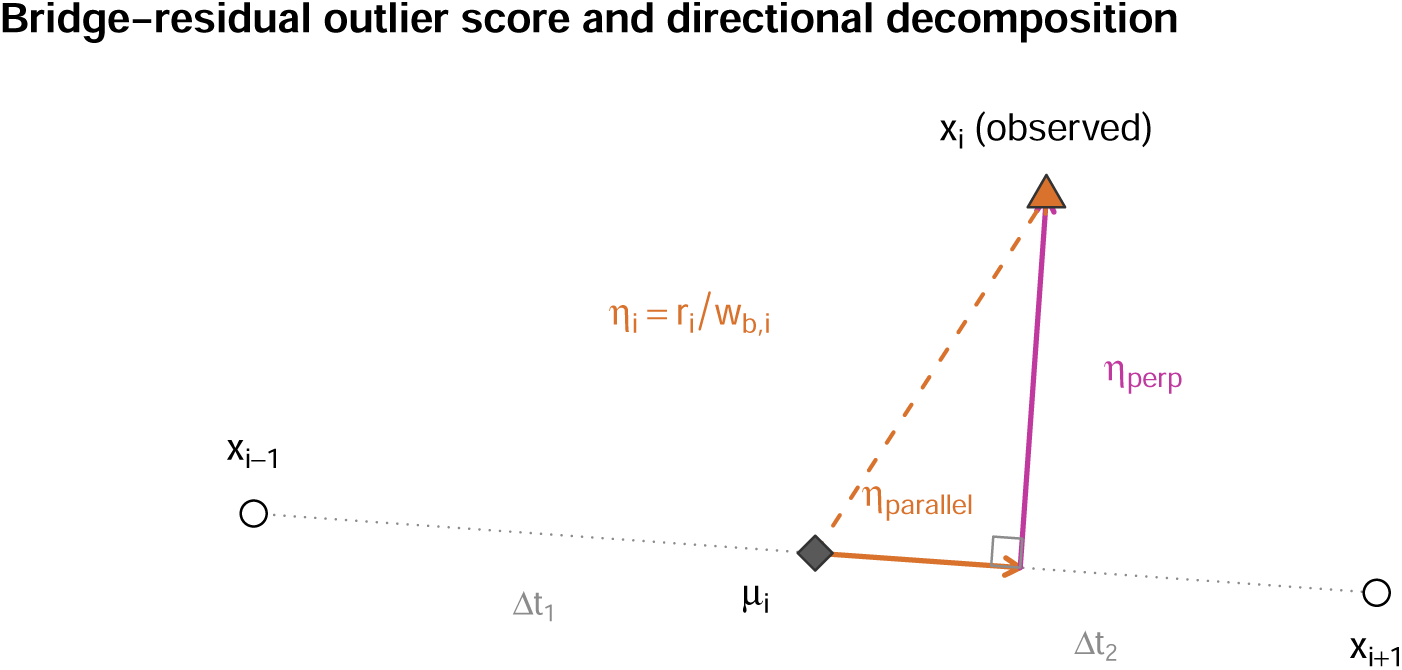
Bridge-residual outlier score and directional decomposition. Three consecutive locations **x***_i−_*_1_, **x***_i_*, and **x***_i_*_+1_ with timestamps defining gaps Δ*t*_1_ and Δ*t*_2_. The bridge mean ***µ****_i_* (diamond) is the time-weighted midpoint between the neighbours. T<u>he residual</u> **r***_i_* = **x***_i_* ***µ****_i_* (dashed arrow), normalised by the bridge width 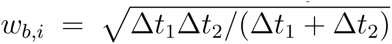 yields the scalar score *η_i_* = *r_i_/w_b,i_* used by method = "isotropic". The directional variant (method = "directional") decomposes the resid- ual into components parallel (*η_‖_*) and perpendicular (*η_⊥_*) to the local travel axis, enabling error-morphology classification. The default, method = "combined", thresholds both *η* and *η_⊥_* independently and flags any location caught by either.

The probability-based method of Section S1.1 (see Eqs. 1–4) scores each location using quantities (speed, angular velocity, and their consecutive changes), that are inherently about *transitions* between locations. It is at its best for spike-shaped outliers, where a single location produces extreme transitions on both sides. It is at its weakest for blocks of outliers, where multiple consecutive locations sit together in the wrong place: within the block, the transitions between neighbours look ordinary, and only the boundaries of the block carry a detectable metric signal. The result is that a drift cluster of twenty locations typically produces at most two transition-extreme points (the entry and the exit), while the remaining eighteen blend into the bulk.

A geometric test can reach these interior points directly. For each location *i* with temporal neighbours *i −* 1 and *i* + 1, the Brownian bridge provides an expected position conditional on the neighbours and the elapsed times: the time-weighted mean

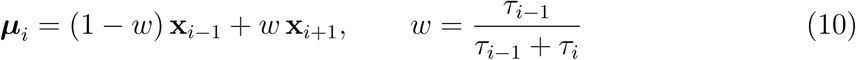

where *τ_i−_*_1_ = *t_i_ − t_i−_*_1_ and *τ_i_* = *t_i_*_+1_ *− t_i_* are the gaps to the preceding and following location respectively. A standard Brownian motion between **x***_i−_*_1_ and **x***_i_*_+1_ has variance at the bridge mean proportional to *τ_i−_*_1_*τ_i_/*(*τ_i−_*_1_ + *τ_i_*), whose square root defines the bridge width

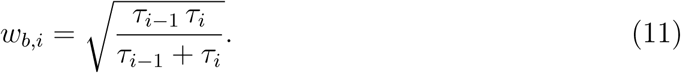

The gap-aware geometric score is the residual normalised by the bridge width:

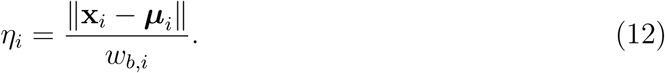

*η_i_* has units of 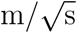 and is, under a Brownian process with any fixed diffusion coefficient, distributed as that coefficient times a folded bivariate-normal magnitude. Its value does not depend on any locally estimated variance. This matters: a variance estimate at location *i*, whether from a dynamic Brownian-bridge model (Kranstauber *et al*., 2012) or from a simpler local fit, is inflated by the very outlier the score is trying to detect, so any score of the form residual-over-local-standard-deviation systematically undershoots at the point where it should overshoot. The bridge-width normalisation (Eq. 11) avoids this leverage by depending only on the timestamps, which cannot be corrupted by the position outlier.

The outlier score *η_i_* is thresholded with the same self-determining detectors applied to log *P*_joint_ in Section S1.3. The default is the density-valley detector applied to *−* log *η* (equivalently the upper tail of log *η* plotted in Figure 3): outliers are declared only when the density of the score distribution contains a real dip between bulk and tail. On clean tracks the distribution of log *η* is unimodal with no such dip, and the detector returns no outliers; the detector is thus a no-op on the clean tracks tested here, which are regularly sampled or duty-cycled with stochastic dropouts. That scope is not incidental. The score is gap-normalised under a Brownian null, and movement that is not Brownian leaves a residual gap dependence, so on a strictly two-valued day–night schedule, long gaps and short gaps with nothing in between, the detector can fire on the sampling schedule rather than on the movement. Neighbour dedup (a greedy peak-picking step that keeps only the largest *η* within each run of consecutively flagged indices) suppresses the smearing artefact by which an outlier at *i* distorts the bridges centred on *i −* 1 and *i* + 1.

#### Directional decomposition (dBGB variant)

Decomposing the bridge residual into components parallel and perpendicular to the local travel axis turns the same detector into an error-morphology classifier. Let **a**^*_i_* be the unit vector from ***µ****_i_* towards **x***_i_*_+1_; the residual **r***_i_* = **x***_i_ −* ***µ****_i_* decomposes into

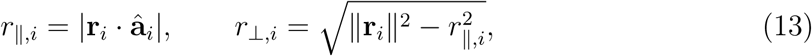

and the gap-normalised directional scores are *η_‖,i_* = *r_‖,i_/w_b,i_* and *η_⊥,i_* = *r_⊥,i_/w_b,i_*. The naming follows the dynamic bivariate Gaussian bridge model from which it is adapted (Kranstauber *et al*., 2014), though the residual score itself does not require the dBGB variance estimate; bridge width alone serves as denominator, preserving the leverage- immunity of Eq. 12. The directional variant flags on *η_⊥_* specifically. The interpretation is that errors which drift perpendicular to legitimate movement (multipath reflections, GNSS spoofing clusters displaced off the flight path, tag-to-tag handoffs), show up cleanly on the orthogonal axis, while along-track behavioural variation (bursts of fast flight, stops and accelerations) falls on the parallel axis and is not flagged. The full pair (*η_‖_, η_⊥_*) forms a two-dimensional diagnostic space in which a flagged point’s location reveals its error morphology: perpendicular-dominant, along-track-dominant, or magnitude-extreme on both axes. The framework also works in reverse: persistent perpendicular-drift flags concentrated at one location are consistent with location-dependent multipath (the colony-halo case in CPF_E) and could be used to down-weight that location even when no single location is an outlier on a scalar test. On the synthetic cohort this morphology mapping holds by construction.

#### Combined detection (default)

Because *|r_⊥_| ≤_‖_***r _‖_** , the scalar score *η* is an upper bound on *η_⊥_*, and flagging on *η* alone includes all points *η_⊥_* would flag plus the along-track extremes. The two scores are correlated but not redundant: a location whose residual is dominantly parallel (physiological speed variation, behavioural transitions, or a short burst of rapid movement), inflates *η* but leaves *η_⊥_* near baseline. Conversely, a location whose residual is dominantly perpendicular (multipath, spoofing drift, tag-to-tag handoff), can sit at a modest *η* even when its *η_⊥_* is extreme, because the large parallel component dilutes the scalar magnitude. Each score therefore has access to a class of outlier the other cannot reach without tuning.

The default software behaviour composes these tests: the self-determining threshold is applied independently to *−* log *η* and to *−* log *η_⊥_*, and a location is flagged if either test trips. On synthetic ground truth (Section 3.1) this combined detector Pareto-dominates both single-score variants: the scalar path recovers strong-omnidirectional outliers (CPF_D’s sustained spoofing block) and the orthogonal path recovers perpendicular- leverage outliers (CPF_E’s colony halo) that the scalar magnitude dilutes. The two paths remain exposed as method = "isotropic" and method = "directional" on mt_flag_outliers_bridge() for studies that deliberately restrict the detection to one error morphology, or for comparability with work that reports one score in isolation. State-aware variance-estimating siblings mt_flag_outliers_dbbmm() and mt_flag_outliers_dbgb() are additionally available for users who want to substitute the full dBBMM/dBGB variance for the bridge-width normalisation and apply a Bonferroni-controlled multi-channel test on the directional decomposition, at the cost of relaxing the leverage-immunity guarantee of Eq. 11.

### S1.5 The detour detector

The bridge residual gains its leverage immunity from temporal geometry, but its sensitivity to single-location excursions degrades as the sampling interval grows, because the bridge width widens with the gaps it is built from. At hourly GPS sampling, an out-and-back excursion to a colony halo can leave each bridge residual within the expected diffusion envelope while still being a clear geometric impossibility.

The detour ratio addresses this regime. For each location *i* and a window radius *k*, define the path length and the chord displacement on the local window:

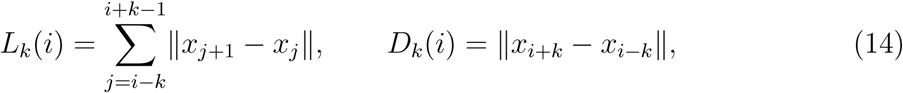

and the dimensionless detour ratio

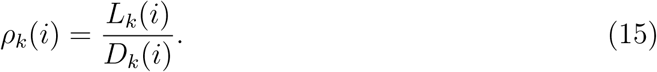

By construction *ρ_k_ ≥* 1 with equality on a straight line, and on a smooth track *ρ_k_* remains close to unity for any reasonable *k*. A symmetric out-and-back excursion at location *i* inflates *L_k_* without commensurate change in *D_k_*, sending *ρ_k_*(*i*) *→ ∞*. A location is flagged if *ρ_k_*(*i*) *> ρ^∗^* for any *k* in a small set of window radii (default *k* = 1).

Two properties of *ρ_k_* matter for its scope of validity. It is time-insensitive: the ratio depends only on consecutive coordinates, not on the time gaps between them. It is invariant to spatial scale: as a dimensionless ratio of two lengths, its threshold needs no calibration to the animal’s typical step, which is why it is the one detector that can carry a fixed constant at all. Its operating range is set instead by movement persistence. For roughly equal steps *ρ*_1_ = 1*/|*cos(*φ/*2)*|* with *φ* the turning angle, so the null depends on the turn-angle distribution: under persistent movement a fixed *ρ^*^* = 8 fires on essentially nothing, while under uniform turning it would fire on 8% of uncontaminated locations.

Tracking data sampled faster than the velocity decorrelation time is persistent, which is the regime the detector is for and the regime the cohort occupies. Where the bridge detector loses sensitivity at sparse sampling, the detour detector does not, which is what the pairing buys. The two are separate constructions witnessing the same fact, that a position is geometrically incompatible with its temporal neighbourhood, but at the default window they read the same three locations, so their agreement is corroboration rather than independent confirmation. When both fire, they constitute geometric consensus, sufficient evidence for outlier status without any kinematic confirmation.

The default threshold differs between the two call sites. Inside the cleaning cascade of Section 2, mt_clean_track() passes *ρ^∗^* = 8, chosen conservatively rather than fitted. The value is not the smallest that preserves the clean reference: at move2utils v0.4.5 the cascade returns zero flags on CPF_B at every *ρ^∗^* from 3 to 8, so the cohort does not discriminate within that range and the constant should be read as a conservative choice rather than a calibrated one. The standalone mt_flag_outliers_detour() call defaultsto *ρ^∗^* = 5, the more sensitive setting appropriate when no second detector gates the flag. Both values are HEURISTIC; the plausible range documented for the detour detector is *ρ^∗^ ∈* [3, 10].

### S1.6 The speed cap

The three detectors above examine the location’s relationship to its temporal neighbour- hood; they say nothing about whether the step out of, or into, that location lies within the species’ physiological capacity. A complementary detector operates at the step level.

Let *v_i_* =_‖_ *x_i_*_+1_ *− x_i‖_ /*(*t_i_*_+1_ *− t_i_*) be the step speed entering location *i* + 1. A location is speed-flagged if its incoming or outgoing step exceeds a cap *c*:

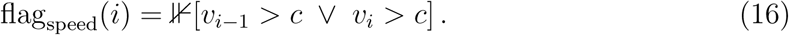

The cap *c* admits four threshold modes (parameter threshold_type):

1. auto: a data-driven cap recovered from the empirical *v* distribution by entropy- valley search. Where that search finds no valley, a broken-stick fallback is used, and it is that fallback which is gated by Hartigan’s dip test of unimodality (Hartigan & Hartigan, 1985) at *α* = 0.05; the entropy-valley candidate is not dip-tested. A candidate cap is accepted only if (a) the implied flag fraction lies below an internal safety guard (5% of locations), and (b) the cap does not fall between two substantive activity modes (an internal mode-position guard). The data-driven cap also fires a warning if it exceeds 55 m/s, a deliberately loose cross-taxon ceiling on *sustained* speed: nothing living holds such a speed over the interval separating two locations, so a step implying one is not the animal’s movement.
2. entropy: the entropy-valley boundary alone (without the dip-test gate). Less con- servative.
3. gap: the broken-stick gap detector applied to sorted *v* (MacArthur, 1957). Detects extreme tails separated from the bulk by an unusually large gap.
4. hard: the user supplies a numerical cap directly. The recommended route to a hard cap is the allometric maximum-speed scaling law of Hirt et al. (Hirt *et al*., 2017), exposed by the package’s v_phys_estimate(mass, mode) function. For body mass *M* in kilograms and mode *∈* {running, swimming, flying}, the realised maximumspeed is

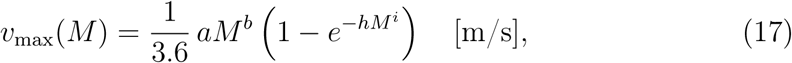

with mode-specific parameters (*a, b, h, i*) taken from (Hirt *et al*., 2017, Supplementary Table 4). Hirt et al. report parameters such that the right-hand side evaluates to *v*_max_ in km/h; the leading 1*/*3.6 converts the result to the SI step-speed units used throughout this paper. The first factor is the power-law ceiling that would be reached if acceleration time were unlimited (with exponent *b* of 0.24 for flying, 0.26 for running and 0.36 for swimming) and the saturating term captures the finite anaerobic energy budget that limits the largest animals to a fraction of that ceiling, making the law accurate across many orders of magnitude in body mass. The law is built on maximum *anaerobic* (burst) capacity, and Hirt et al. explicitly excluded the maxima of normal speeds, including dispersal and migration. That is what makes it the right basis for a coarse impossibility bound rather than a movement model: nothing an animal sustains in ordinary travel comes near it.

**Table 4:** The cascade’s conjunction classes. Each row defines a sufficient condition for flagging location *i*, written over the binary detector firings *b_i_* (bridge), *δ_i_* (detour), *p_i_*(autodifference probability), and *s_i_* (state-relative speed), each 1 when that detector fires and 0 otherwise; and are logical AND and OR. Single-detector firings never flag in isolation: every class requires either two independent geometric witnesses (*b_i_, δ_i_*), a geo- metric witness plus a kinematic witness (*p_i_* or *s_i_*), three-of-four consensus, or a structural witness from outside the per-location vote (block expansion, pre-peel). The *Methodological reading column paraphrases each predicate.*

| Class | Predicate | Methodological reading |
| --- | --- | --- |
| *consensus* | $b_i + \delta_i + p_i + s_i \geq 3$ | Three of four detectors agree |
| *geometric_spike* | $b_i \wedge \delta_i$ | Two independent geometric witnesses |
| *state_anomaly* | $(b_i \vee \delta_i) \wedge s_i$ | Geometric witness + state-relative speed |
| *kinematic_confluence* | $(b_i \vee \delta_i) \wedge p_i$ | Geometric witness + autodifference probability |
| *block* | block expansion fired | Outside the dominant along-track segment (Section S1.8) |
| *physiological* | pre-peel fired | Step exceeded a hard physiological cap (Section S1.6) |

The 55 m/s ceiling is a structural sanity bound, not an empirical cutoff, and what sets it is what living animals can sustain rather than any fit. A stooping falcon exceeds it for seconds; nothing alive holds it over the minutes to hours between two locations, so a step speed above it is not the animal’s movement but a positional error or a tag that is no longer on the animal. The check is advisory: it emits a message and never alters the cap. Supplying v_phys_estimate(mass, mode) sharpens that universal envelope into a per-species ceiling for the cost of two metadata fields, and because it derives from physiology rather than the track it cannot be inflated by the very outliers it catches.

The speed cap appears in the cascade of Section 2 in two distinct scopes. In the *pre-peel* stage, an external physiological cap (mode hard, supplied via (mass, mode) or a manual value) is iteratively applied to remove biologically impossible steps before any other detector sees the track. In the *conjunction* stage, a data-driven cap (mode auto) operates state-relative to the per-segment distribution and contributes one of the four detectors that the cascade combines. Distinguishing these scopes is necessary because a physiological cap is a sanity bound that cannot be exceeded, whereas a data-driven cap is a within-distribution diagnostic that varies with the animal’s behavioural state.

### S1.7 The conjunction structure

For each location *i*, let *b_i_*, *δ_i_*, *p_i_*, *s_i_ ∈ {*0, 1*}* denote the firings of the bridge, detour, probability, and state-relative speed detectors. The class-aware conjunction flags *i* as an outlier if any one of the classes in Table 4 fires; the error_class returned for *i* is the highest-priority class that fired (in decreasing priority: *physiological*, *block*, *consensus*, *geometric_spike*, *state_anomaly*, *kinematic_confluence*).

Each class is a logical conjunction whose constituent detectors operate within non-overlapping scopes of validity. Geometric witnesses (bridge, detour) are valid regardless of behavioural state; kinematic witnesses (state-relative speed, autodifference probability) are valid only relative to the local distribution. Combining a geometric and a kinematic witness across these scopes yields a witness pair where neither weakness gates the other’s strength. This is the principle that distinguishes the cascade from a majority vote, in which the weakest detector at sparse sampling can effectively veto the strongest’s correct firing.

The default rule, evidence_corroborated, generalises this conjunction from a Boolean test to an accumulation of evidence. Each detector contributes a signed evidence score *ℓ_k_* (its own score at location *i* minus its own data-driven flag boundary, a per-detector standardised score rather than a likelihood ratio under a shared generative model), so that *ℓ_k_ >* 0 essentially when detector *k* fires and grows with how far past its threshold the location lies. The relation is not an identity, because the boundary is the least extreme score among the locations that detector has already flagged: the location that sets it scores exactly zero, and a location the detector declined for some other reason can score above it. That is also why the corroboration count behaves like a vote rather than like a sum. Only two of the four are surprisals: the probability detector contributes *−* log *p_i_* and the bridge <u>^1^</u> *η*^2^, while the detour and the speed cap contribute the log-transformed statistics log *ρ_i_* and log *v_i_*.

These native magnitudes are incommensurable, since a bridge residual can reach the millions while a log-ratio sits near unity, so each is calibrated to a common, bounded scale, *C* tanh(*ℓ_k_/*MAD*_k_*), with the median absolute deviation taken over the track, so that no detector can dominate by units alone. A location is flagged when the summed calibrated evidence *E_i_* = Σ*_k_ C* tanh(*ℓ_k_/*MAD*_k_*) is positive *and* either at least two detectors contribute positively, or the detour detector, the one geometric witness an out-and-back cannot fake, is saturated, its calibrated evidence at least two MADs beyond its own flag boundary (exceeding *C* tanh 2). The first clause is the unsupervised regime boundary between the homogeneous inlier mass and the conspicuous tail; the second restores the corroboration discipline of the conjunction while still letting a single overwhelming geometric witness carry.

The combined evidence *E_i_* is returned as a column, giving the user a continuous suspicion score and a single lever on sensitivity. The rule’s own thresholds are properties of the track, with four exceptions that are fixed: the detour ratio’s *ρ^*^*, the saturation level at which a single detector may carry a flag alone, the cut on the summed evidence, and the calibration ceiling *C*. The last two are fixed without being tuned. The cut sits at zero because each *ℓ_k_* is already centred on its own detector’s flag boundary, so zero *is* that boundary rather than a chosen value; and *C* cannot change any flag at that cut, since it is a common factor of the sum, the corroboration count is taken on the uncalibrated scores, and the saturation test compares *C* tanh(*ℓ/*MAD) against *C* tanh 2: it cancels from all three clauses and scales only the reported score.

The class-aware conjunction remains available as consensus = "class_aware", with which the default coincides on the easy and the clean cases (Supplement S7.1). Sev- eral further rules ship for users who want a different precision-vs-recall trade-off without rebuilding the cascade: "strict" ((bridge *∨* detour) *∧* (prob *∨* speed)), the highest-precision rule; "majority" (any 2 of 4 detectors agree); "speed_trusted" (speed alone is sufficient, on the grounds that its auto mode is already dip-test-validated); "weighted_evidence" (the same evidence accumulation as the default but without the corroboration requirement); and "any" (union of all four, maximum recall, lowest preci- sion); plus "custom" for a user-supplied function over the four detector firings. All are documented in ?mt_flag_consensus and operate on the same per-detector flag columns the cascade emits, so users running detectors outside the cascade can apply them post-hoc.

### S1.8 Block expansion

Per-location detection structurally cannot resolve a coherent train of outliers whose internal geometry is locally smooth.

After the per-location conjunction has labelled candidate outliers, the cascade constructs a graph *G* = (*V, E*) on the kept locations. An edge joins two *temporally consecutive* kept locations if the step between them would satisfy the physiological cap: that is, ‖ *x_i_*_+1_ *−x_i_‖ /*(*t_i_*_+1_ *−t_i_*) *≤ c*_phys_. The graph is therefore a path, and its components are maximal runs of kept locations uninterrupted by an over-cap step; the step is a segmentation along the track rather than an all-pairs construction. That restriction is consequential: a block lying mid-track cannot be isolated without severing the legitimate trajectory, which is why the dominance gate below declines it. One clause of the implementation is not in that expression and is load-bearing rather than incidental: where flagged locations have been removed *between* two kept locations, the lag used for the edge test is capped at the track’s median consecutive sampling interval. Without it, peeling a block’s seam locations would widen the kept-to-kept gap until the implied across-seam speed fell below the cap and the block re-joined the main component, so a supplied ceiling could reduce recovery rather than improve it. Genuine single-step gaps, including real missing-data gaps, keep their true lag. The graph is then decomposed into *k* connected components*{C*_1_*, C*_2_*, . . . , C_k_}* ordered by size, with *|C*_1_*| ≥ |C*_2_*| ≥ · · · ≥ |C_k_|*.

Block expansion is gated by two conditions, both of which must hold before any non- largest component is flagged. First, the largest component must dominate the partition:

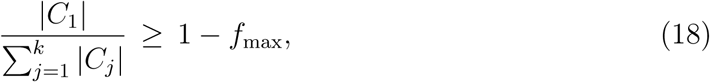

where *f*_max_ is an internal fraction (bound to max_flag_fraction, default 0.2, so the largest component must hold at least 80 % of the kept locations). This guards against a too-tight cap: a continuous trajectory carved into many similarly-sized fragments fails this test and expansion is declined. Second, when *k ≥* 4, a clean log-gap must separate the largest component from the bulk:

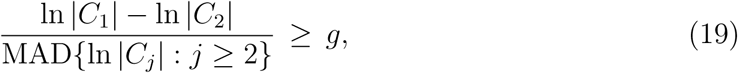

where *g* is an internal gap factor (default *g* = 3; the value reuses the broken-stick three-sigma convention applied elsewhere in the package). The numerator is the log-gap between the largest and second-largest component; the denominator is the median absolute deviation of the log-sizes of all non-largest components, used as a robust dispersion baseline. This mirrors the broken-stick / log-gap-above-bulk reasoning the package applies to per-step speeds; here it is applied to the upper tail of the log-component-size distribution. When *k ∈ {*2, 3*}*, only Eq. 18 is required because there are too few components to estimate a stable bulk dispersion. When the gate passes, every location in a non-largest component is flagged and receives error_class = "block" in priority order over the per-location classes.

The graph construction depends on a meaningful *c*_phys_. When the user supplies (mass, mode), *c*_phys_ is the Hirt et al. allometric *v*_max_ (Eq. 17); otherwise the cascade uses the data-driven auto cap as a fallback, with the safety guard documented in Section S1.6. Two things limit what block expansion can do: it cuts along the track, so a block interleaved with real locations never forms a segment of its own, and it runs last, on whatever the detectors have left.

### S1.9 State-conditional cleaning and the global rescue pass

Real GPS tracks rarely sample a single behavioural distribution. A migrating bird’s velocity at the colony shares no quantile of the velocity distribution with its velocity in flight; a terrestrial mammal’s resting jitter shares no quantile with its commuting steps. Pooling these into a single distribution dilutes the threshold that any kinematic detector tries to recover, and introduces a regime in which the right cap for one state is wrong for another.

The package’s mt_clean_track(state = *·*) call accepts either a column name in the input move2 object or a per-location vector of state labels. The cascade partitions each track into contiguous runs of constant state and runs the kinematic detectors per segment. Per-segment thresholds adapt to each state’s distribution, sharpening sensitivity within each segment and preventing one state’s bulk from setting the threshold for another’s tail. Per-segment dispatch sharpens the kinematic detectors by construction, but it also changes what the geometric detectors see: and there it does damage. The bridge residual at location *i* measures geometric impossibility relative to its temporal neighbourhood; the detour ratio is dimensionless. Neither concept depends on which behavioural state the animal is in. If the per-segment partition splits a colony halo across two states (typical for HMM segmentation, where a brief out-and-back at the colony is mis-assigned to migrate), the bridge threshold within the migrate segment is inflated by legitimate flight residuals and the geometric impossibility goes undetected.

To prevent this, the cascade runs a *global rescue pass* in addition to the per-segment passes. Concretely, for each per-location detector *D ∈ {*bridge, detour, prob, speed*}*, let *D*_seg_(*i*) *∈ {*0, 1*}* be the firing of *D* when run on the segment containing location *i*, and let *D*_glb_(*i*) *∈ {*0, 1*}* be its firing when run on the full track. The cascade defines the effective firing as the union of the two:

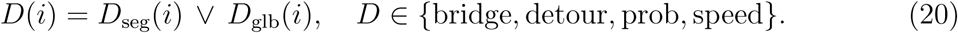

The geometric witnesses (bridge, detour) benefit most from the global pass, since their notion of impossibility is state-independent and a state-segmented threshold can only dilute it; the kinematic witnesses (prob, speed) contribute additional flags for steps that are anomalous against the full-track distribution even when they sit within their own state’s distribution. The conjunction classes of Section S1.7 are then evaluated on the combined firings.

### S1.10 Transition handling

State labels assigned by an HMM, a speed threshold, or any other segmentation procedure are themselves uncertain in the immediate neighbourhood of a state change. Within *±k* locations of any detected transition (default *k* = transition_buffer = 1), the cascade demotes any flag whose error class is state-dependent: *state_anomaly* and *kine- matic_confluence* are re-labelled *state_transition_buffered* and the location is kept. The state-independent error classes (*geometric_spike*, *consensus*, *block*, *physiological* ) continue to flag inside the transition buffer, since their predicates do not depend on the state assignment being correct.

### S1.11 Adapting the cascade to real data

This subsection documents the first-class arguments summarised in Section 4.

#### Pooling across deployments and individuals (pool_by)

When the same animal carries successive tags or a study spans a behaviourally homogeneous population, the per-track defaults under-use the data: each track’s detector thresholds are fitted to that track alone, even when many tracks could share a more precise threshold. Providing a single column to pool_by = <COLUMN> (typically individual_id, study_id, or species) runs a post-cascade sweep that refits the bridge, detour, and speed cap detectors once on each pool group and unions the pool-added flags (error_class = "pool") into the per-track result. The cascade itself stays per-track and is byte-identical to the default in single-individual workflows, so pool_by costs nothing on studies that do not need it. The single-column form uses one group for both the threshold-fitting distribution and the flag union; when these roles differ (for example, drawing threshold strength from a whole cohort while keeping the union scoped to the individual), pool_by accepts c(outer, inner), the *outer* level supplying the fitting distribution and the *inner* bounding the union, under strict nesting. Each call assumes a single error regime, so tracks interleaving sensors of very different precision (e.g. metre-scale GPS and kilometre-scale geolocation) should be split by sensor before pooling and re-merged after.

#### Behavioural states and the global rescue pass (state)

Many tracks sample qual- itatively different behavioural modes (resting versus migrating, perching versus flying), whose kinematics share almost no quantile of the speed or turn-angle distribution, so pooling them forces the kinematic detectors to over-flag the slow mode or under-flag the fast one. A per-location state vector (from a fitted HMM (McClintock & Michelot, 2018; Michelot *et al*., 2016), a speed-threshold split, or manual annotation) runs the kinematic detectors within each contiguous state run, while the state-independent geometric detec- tors (bridge, detour) are additionally run on the full track; their union is the global rescue pass (Section S1.9). A transition_buffer demotes state-dependent flag classes within *±k* locations of a transition (default *k* = 1), where the state label itself is uncertain.

### Other options

location_error supplies a per-location observation-error prior (a nu- meric vector in metres, or "auto" to read Movebank quality columns), by which the bridge inflates its width so locations consistent with reported uncertainty are demoted. pre_peel_aux = "detectors" switches the pre-peel stage from flagging both endpoints of an over-cap step to flagging only the higher-scoring one, useful on tracks dominated by isolated spikes. persistence_filter = "class_aware" demotes low-persistence flags in the state_anomaly and consensus classes. Five further NULL-defaulted overrides forward non-default values into the detector calls for replication or diagnostics; all defaults and plausible ranges are catalogued in Supplement S3.

## S2 The cascade in pseudocode

The listing below is the whole procedure in one place, for a reader who wants the order of operations rather than the derivations. Section and equation references point to the subsections above.

Algorithm 1 mt_clean_track cleaning cascade

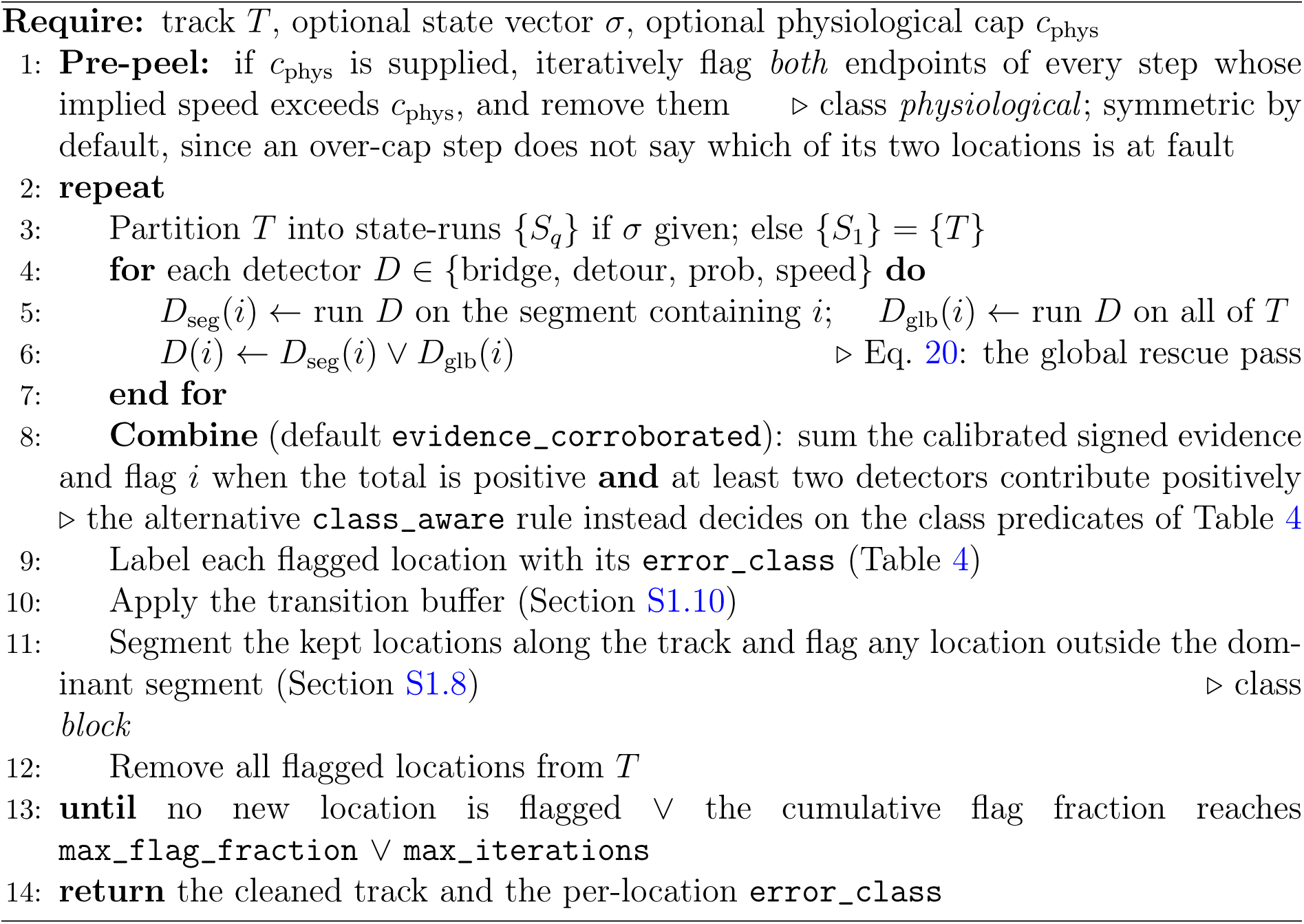

Two properties of the loop are worth reading off the listing. Flags accumulate *across* iterations, so a coherent train of outliers is peeled from its seams over successive passes rather than recognised at once: on the cohort’s spoof track the recovery takes 28 iterations, one truth location per pass but for a single pass that took three. And the block step runs last, on the locations the per-location layer has left, so it only ever sees what that layer did not resolve.

## S3 HEURISTICS catalogue

The package maintains an explicit accounting of every non-derived constant in the outlier-detection pipeline, following the design principle of “no silent magic.” Each constant is classified as **DERIVED** (traces to a published equation, theoretical bound, or measured constant), **HEURISTIC** (a defensible round number with a stated rationale and plausible range; the choice within that range is convention), or **EMPIRICALLY TUNED** (selected to fire on a specific benchmark case, and flagged for replacement by a structurally-grounded alternative). Tables 5, 6 and 7 give the constants of each class in turn. They follow the outlier-detection entries of the package’s living catalogue (move2utils/HEURISTICS.md); utilities outside the scope of this paper (e.g. mt_corridor) are catalogued there but omitted here. The catalogue is a living document, and the two can differ in how a constant is justified rather than in the value it takes; where they do, the basis given here is the one this paper stands behind. None of these constants is set by the user through mt_clean_track()’s default call; the “plausible range” column states the convention band, not a tuned optimum.

**Table 5:**
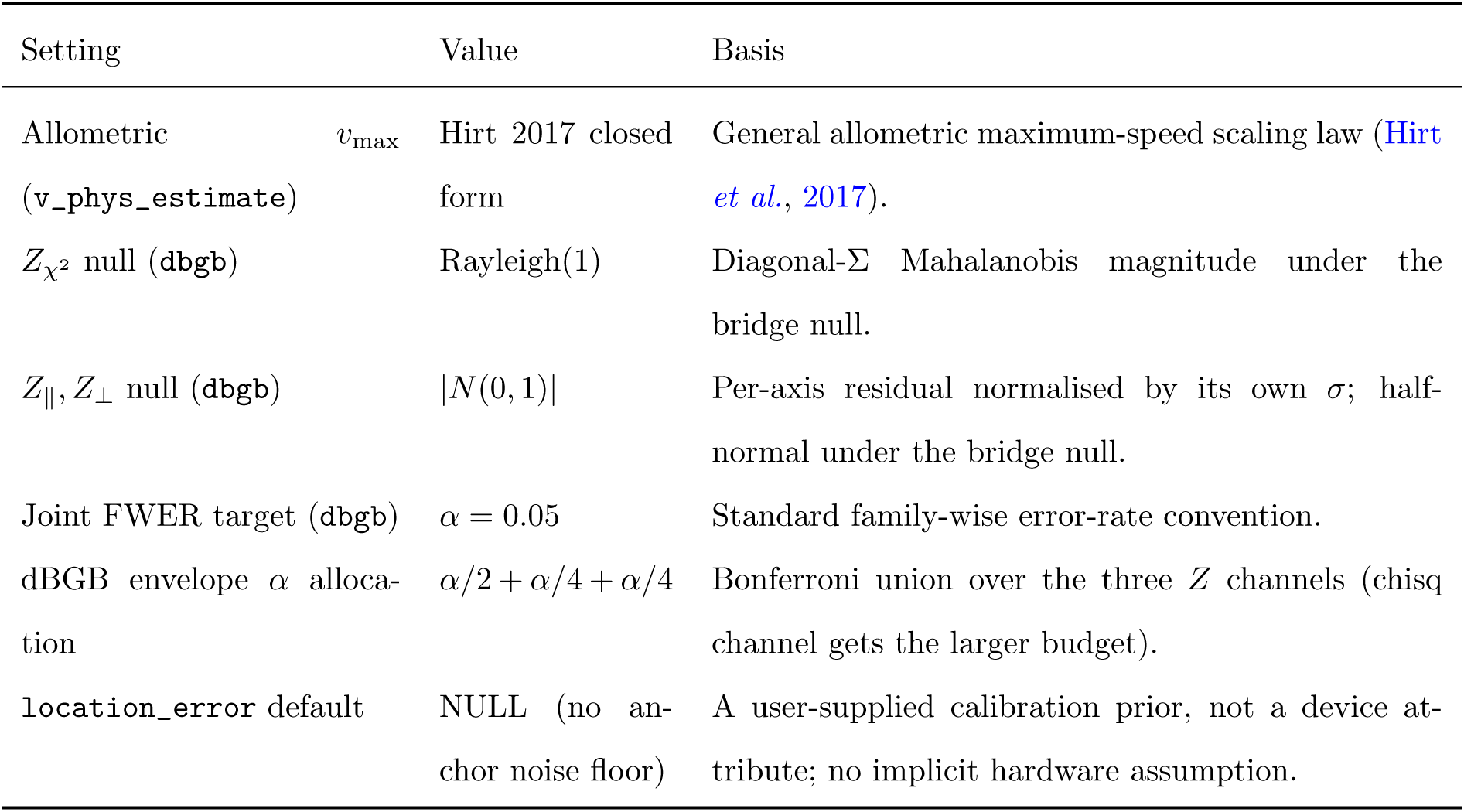
DERIVED settings: traceable to theory or measurement.

**Table 6:**
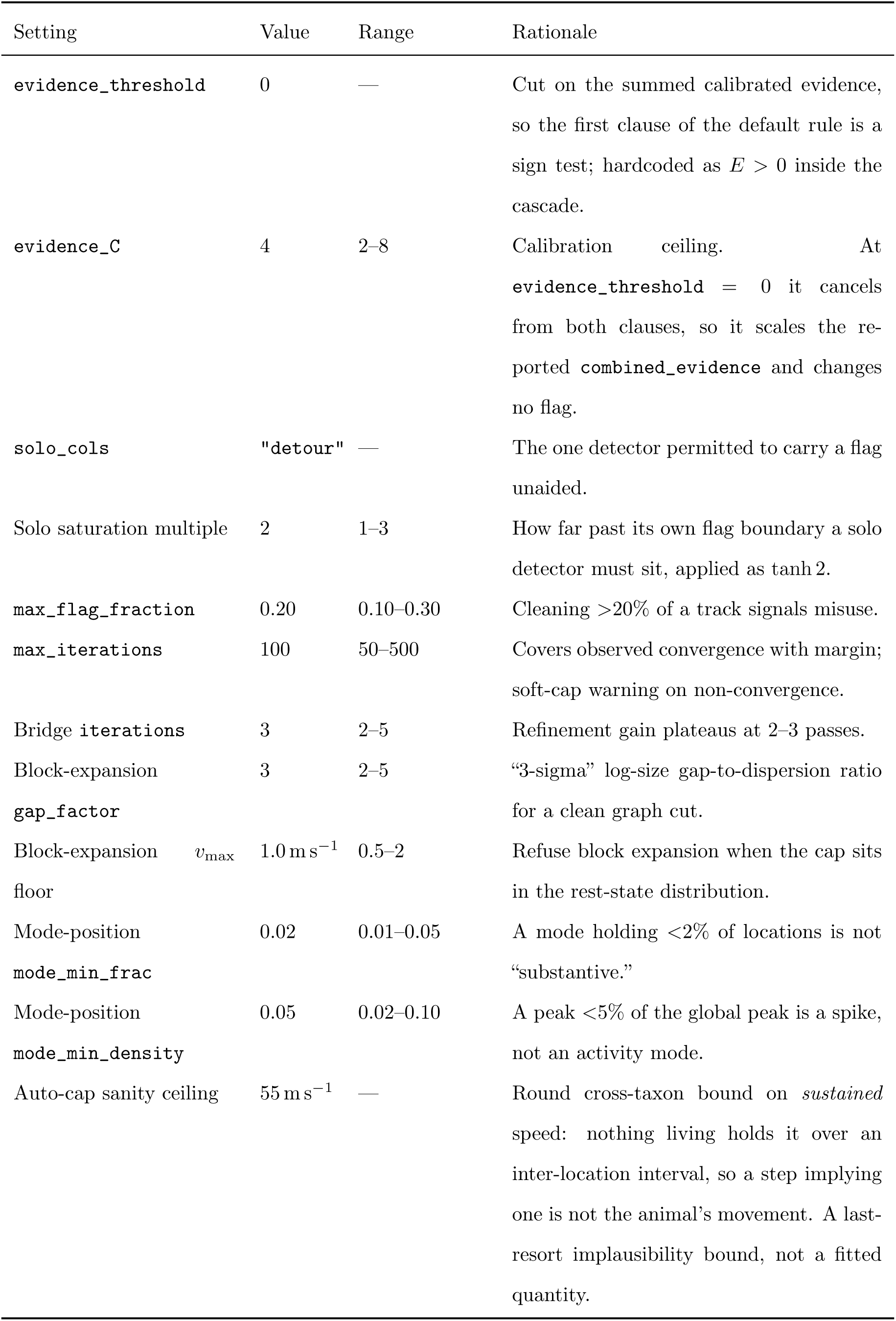
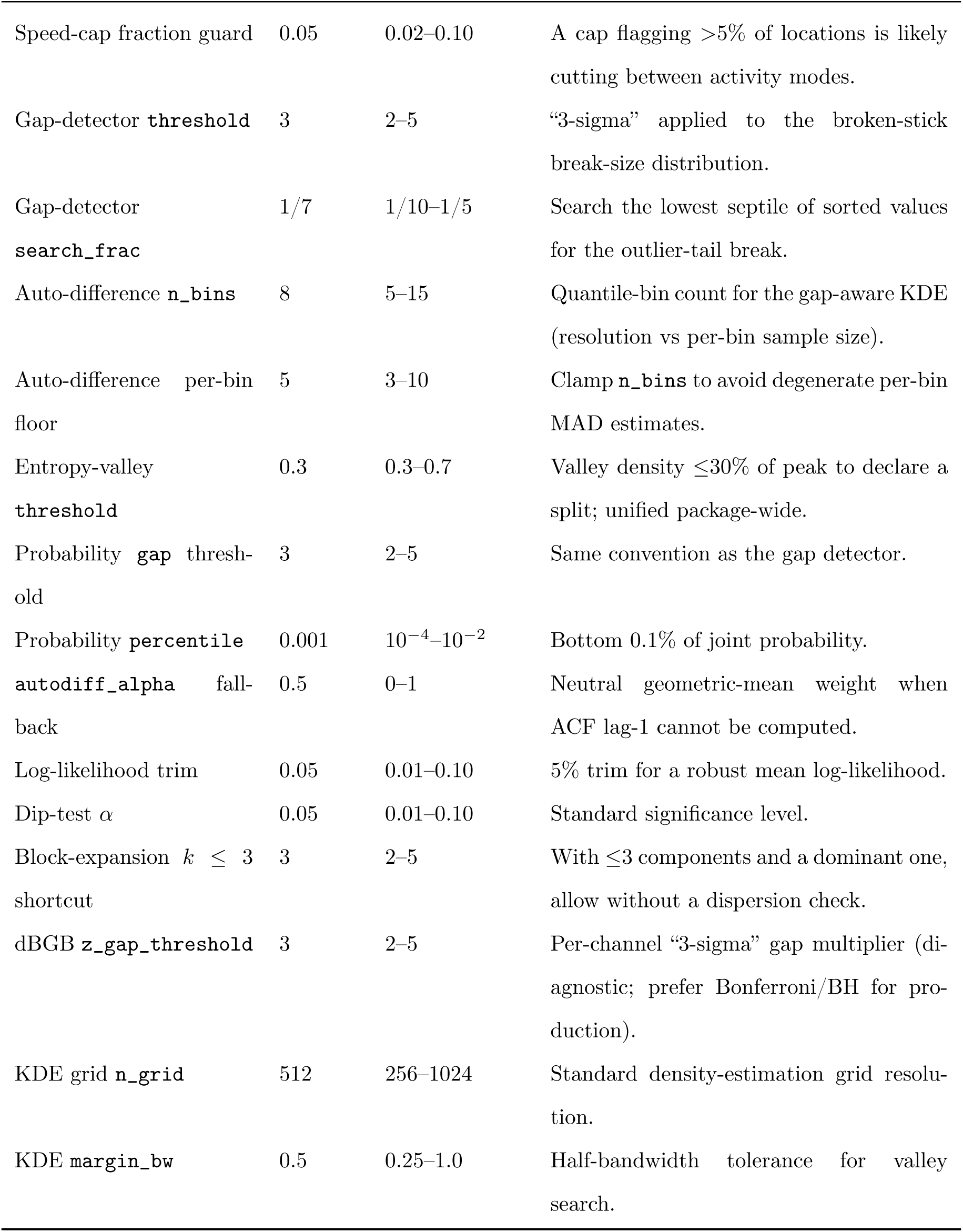
HEURISTIC settings: defensible round numbers with a stated rationale and convention band.

**Table 7:** EMPIRICALLY TUNED settings: selected against a benchmark case and flagged for replacement by a structural alternative.

| Setting | Value | What it was tuned to / replacement plan |
| --- | --- | --- |
| `detour_threshold` | 8 (cascade) / 5 (standalone) | Dual default: the cascade conjunction lets detour stay strict (8) without losing recall; the standalone favours solo sensitivity (5). Calibrate the stability range empirically. |
| `transition_buffer` | 1 | Fixes either side of a state change in which state-dependent classes are demoted to `state_transition_buffered`; hand-set conservatively. Calibrate across a multistate cohort. |

## S4 Methods comparison and the block-expansion demonstration

### S4.1 Detection metrics

Every comparison in this supplement and in the main text scores one tool on one track by counting its flags against that track’s known outlier set. A true positive (TP) is an injected outlier the tool flags, a false positive (FP) a legitimate location it flags, and a false negative (FN) an injected outlier it leaves in place. From those three counts,

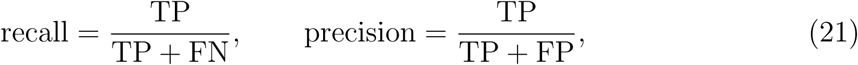

so recall is the share of the outliers present that a tool found, and precision the share of its flags that were outliers. Neither orders two tools on its own: a tool that flags every location has perfect recall, and one that flags a single certain outlier has perfect precision.

Their harmonic mean,

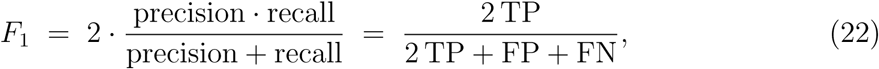

is high only when both are high, and reaches 1 when a tool flags every outlier and nothing else. It is undefined where a track carries no outliers and a tool flags none, TP, FP and FN then all being zero; CPF_B is that case for every tool.

### S4.2 Synthetic boundary-spoof demonstration of block expansion

The graph-based block-expansion step (Section 2) engages whenever a physiological speed ceiling is available, found automatically from a clear gap that a coherent block produces through the impossible transitions at its boundary, or supplied via mass/mode. The syn- thetic cohort of Section 3.1 does not exercise it: CPF_D’s injected block sits mid-track, so cutting the kept-location graph at its impossible boundary transitions severs the legit- imate trajectory into two comparable halves and the dominance gate correctly declines (no component holds the required majority of locations), while the complementary detec- tors and their consensus already recover the block in full. Block expansion is therefore scoped to the case in which the trajectory offers no evidence either way: a *sustained, boundary-anchored* block, where one seam leaves two stretches that cannot witness each other and the majority assumption is the only thing left to decide on. Table 8 reports the demonstration on such a track.

**Table 8:** Block-expansion demonstration on the synthetic boundary-spoof track (150- location block at the track boundary). The cascade recovers the entire block through the graph-based block-expansion step (the block error_class) whether the connectivity ceiling is found automatically (zero parameters) or supplied via mass/mode; a per-location speed cap recovers none of the block interior. Generated by scripts/block_demo.R

| Tool | flagged | TP | FP | recall | precision | $F_1$ |
| --- | --- | --- | --- | --- | --- | --- |
| `mt_clean_track` (zero parameters) | 155 | 150 | 5 | 1.000 | 0.968 | 0.984 |
| `mt_clean_track` (`mass/mode`) | 154 | 150 | 4 | 1.000 | 0.974 | 0.987 |
| naïve speed cap ($> 50 \text{ m s}^{-1}$) | 1 | 0 | 1 | 0.000 | 0.000 | 0.000 |

To demonstrate it on data of known provenance, I constructed a purely synthetic track (scripts/block_demo.R): the clean reference track CPF_B (*n* = 3,537) with its final 150 locations overwritten by a coherent spoof block, pinned at a false coordinate roughly 120 km from the legitimate path (a physiologically impossible boundary transition) with tight within-block jitter so the interior looks locally normal. The block sits at the track’s end, so it disconnects as a single small graph component without splitting the legitimate trajectory. Of the 155 flags the cascade raises, 146 carry the block error_class (the block- expansion output), recovering all 150 truth locations with five false positives at *F*_1_ = 0.984. Supplying a physiological cap via mass/mode recovers this block equally well (150*/*150, *F*_1_ = 0.987), so block recovery here does not depend on whether the connectivity ceiling is found automatically or supplied. That does not generalise: on CPF_D a supplied cap costs recall (1.000 *→* 0.900, *F*_1_ 0.645 *→* 0.607), so more information can reduce recovery.

The naïve speed cap flags only the single boundary transition and reaches none of the block interior, the structural blind spot of per-location scoring. This is the mechanism that, in the main text, distinguishes the cascade on coherent block contamination.

### S4.3 The dedicated reflection filter in atlastools

The headline comparison runs atlastools as a speed-rule filter, which is what a general screen requires and is the configuration that returns zero on CPF_D’s block. The package also ships atl_remove_reflections(), a separate procedure aimed squarely at this error morphology, and it belongs here because it recovers the block the speed rule cannot. Gupte *et al*. (2022) identified prolonged spikes as a distinct class of error and implemented a remedy for them, which is the diagnosis this paper starts from.

The procedure is not a per-location test. It anchors on a location whose in- coming speed exceeds reflection_speed_cutoff and whose turning angle exceeds point_angle_cutoff, then discards every location up to the next one whose outgoing speed exceeds the same cutoff. It takes the track-level view a coherent block demands: it finds the two seams and removes what lies between them.

On CPF_D, at its documented defaults of 20 m s*^−^*^1^ and 45*^◦^*, it recovers all 30 injected locations with three false positives, *F*_1_ = 0.952, against the cascade’s 0.645 on that track (Table 9). On the morphology it was written for, the dedicated instrument is the better one. What it costs is generality. Over the four tracks it completes its mean *F*_1_ is 0.275, against 0.899 for the cascade on the same four. On CPF_A and CPF_E it removes most of the track to reach the outliers, and on CPF_C it recovers none of them. On CPF_B and CPF_F it does not return: no location exceeds both cutoffs, so no anchor exists and the procedure stops rather than returning the track unchanged. That is an assumption about its input rather than a defect, and the documentation states it, asking for data that have already been cleaned. The practical consequence is that it cannot be swept over a study unsupervised: mapped across this cohort in order it stops on the second track, and which tracks it may be applied to at all is a question someone has to answer by inspection first.

**Table 9:** atl_remove_reflections() across the synthetic cohort, at its documented defaults (reflection_speed_cutoff = 20 m s*^−^*^1^, point_angle_cutoff = 45*^◦^*), scored against the same truth sets as Table 2. On CPF_D it recovers the block in full and outscores the cascade (0.952 against 0.645). On the two tracks where no location exceeds both cutoffs it raises an error rather than returning the track unchanged, so it cannot be applied without knowing first that a block is present. Regenerated by scripts/verify_atlastools_reflections.R

| Track | flagged | TP | FP | recall | precision | $F_1$ |
| --- | --- | --- | --- | --- | --- | --- |
| CPF_A | 1003 | 21 | 982 | 0.913 | 0.021 | 0.041 |
| CPF_B (clean) |  | no anchor location: error |  |  |  |  |
| CPF_C | 2 | 0 | 2 | 0.000 | 0.000 | 0.000 |
| CPF_D (spoof block) | 33 | 30 | 3 | 1.000 | 0.909 | **0.952** |
| CPF_E (colony halo) | 1363 | 78 | 1285 | 0.975 | 0.057 | 0.108 |
| CPF_F (multi-state) |  | no anchor location: error |  |  |  |  |

Where the error morphology is known in advance, a filter built for it will do better than a general screen, and on this block it does. The cascade reaches the same block from one call that was given nothing, on a cohort where four of the six tracks carry no block at all and two of them would have stopped this filter outright.

### S4.4 Full per-track, per-tool comparison

Table 10 expands the headline comparison of Section 3.2 into per-track precision, recall, and *F*_1_ for every tool. Every row, competitor and cascade alike, comes from one run of scripts/cohort_competitors.R against move2utils v0.4.5. It makes explicit the per-track structure summarised by the cross-cohort mean: the cascade’s clear margin on CPF_D’s spoof block, its near-parity with a favourably tuned SDLfilter on the easy single-mode tracks, and trip::sda’s tendency to under-fire on this cohort: it is the most precise of the four and the least sensitive, flagging fewer locations than there are outliers on three of the five contaminated tracks and never more than one beyond them. The tier-1 arm, in which the cascade is additionally given mass and mode, is included for every track; it is the arm that shows supplying allometric information *reducing* performance, on CPF_A (*F*_1_ 0.958 *→* 0.885) as well as on CPF_D (0.645 *→* 0.607).

**Table 10:** Per-track precision (P), recall (R) and *F*_1_ for every tool on the truth-bearing tracks, with false-positive counts on the clean reference CPF_B. The tools are as defined for Table 2 (Section 3.1); competitors run at *v*_max_ = 1.5 m s*^−^*^1^ and the cascade at zero parameters. From figures/cohort_competitors.csv.

| Track | Tool | flagged | TP | FP | P | R | $F_1$ |
| --- | --- | --- | --- | --- | --- | --- | --- |
| CPF_A | naïve | 31 | 23 | 8 | 0.742 | 1.000 | 0.852 |
|  | atlastools | 22 | 22 | 0 | 1.000 | 0.957 | 0.978 |
|  | trip::sda | 21 | 21 | 0 | 1.000 | 0.913 | 0.955 |
|  | SDLfilter | 24 | 23 | 1 | 0.958 | 1.000 | 0.979 |
|  | mt_clean_track | 25 | 23 | 2 | 0.920 | 1.000 | 0.958 |
|  | + mass/mode | 29 | 23 | 6 | 0.793 | 1.000 | 0.885 |
| CPF_B | naïve | 0 | 0 | 0 | — | — | — |
|  | atlastools | 0 | 0 | 0 | — | — | — |
|  | trip::sda | 0 | 0 | 0 | — | — | — |
|  | SDLfilter | 0 | 0 | 0 | — | — | — |
|  | mt_clean_track | 0 | 0 | 0 | — | — | — |
|  | + mass/mode | 0 | 0 | 0 | — | — | — |
| CPF_C | naïve | 8 | 4 | 4 | 0.500 | 1.000 | 0.667 |
|  | atlastools | 4 | 4 | 0 | 1.000 | 1.000 | 1.000 |
|  | trip::sda | 4 | 4 | 0 | 1.000 | 1.000 | 1.000 |
|  | SDLfilter | 4 | 4 | 0 | 1.000 | 1.000 | 1.000 |
|  | mt_clean_track | 4 | 4 | 0 | 1.000 | 1.000 | 1.000 |
|  | + mass/mode | 4 | 4 | 0 | 1.000 | 1.000 | 1.000 |
| CPF_D | naïve | 2 | 1 | 1 | 0.500 | 0.033 | 0.062 |
|  | atlastools | 0 | 0 | 0 | — | 0.000 | 0.000 |
|  | trip::sda | 1 | 1 | 0 | 1.000 | 0.033 | 0.065 |
|  | SDLfilter | 4 | 3 | 1 | 0.750 | 0.100 | 0.176 |
|  | mt_clean_track | 63 | 30 | 33 | 0.476 | 1.000 | 0.645 |
|  | + mass/mode | 59 | 27 | 32 | 0.458 | 0.900 | 0.607 |
| CPF_E | naïve | 157 | 80 | 77 | 0.510 | 1.000 | 0.675 |
|  | atlastools | 84 | 80 | 4 | 0.952 | 1.000 | 0.976 |
|  | trip::sda | 81 | 80 | 1 | 0.988 | 1.000 | 0.994 |
|  | SDLfilter | 84 | 80 | 4 | 0.952 | 1.000 | 0.976 |
|  | mt_clean_track | 81 | 80 | 1 | 0.988 | 1.000 | 0.994 |
|  | + mass/mode | 81 | 80 | 1 | 0.988 | 1.000 | 0.994 |
| CPF_F | naïve | 4 | 2 | 2 | 0.500 | 0.500 | 0.500 |
|  | atlastools | 2 | 2 | 0 | 1.000 | 0.500 | 0.667 |
|  | trip::sda | 2 | 2 | 0 | 1.000 | 0.500 | 0.667 |
|  | SDLfilter | 5 | 4 | 1 | 0.800 | 1.000 | 0.889 |
|  | mt_clean_track | 7 | 4 | 3 | 0.571 | 1.000 | 0.727 |
|  | + mass/mode | 7 | 4 | 3 | 0.571 | 1.000 | 0.727 |
|  | + state | 7 | 4 | 3 | 0.571 | 1.000 | 0.727 |
|  | + both | 7 | 4 | 3 | 0.571 | 1.000 | 0.727 |

## S5 Robustness of the comparison

Three experiments vary conditions that the synthetic cohort holds fixed: the speed ceiling handed to the competitors, the regularity of the sampling schedule, and the absence of positional error on legitimate locations. Each is reproduced by a script in the analysis deposit (vmax_sensitivity.R, sampling_sensitivity.R and error_sensitivity.R), and every value quoted below and in the main text is read from the CSV that script writes, with the tables generated in the same run so that they cannot drift from it.

### S5.1 The speed ceiling

Table 11 reports mean *F*_1_ over the five contaminated tracks at each ceiling from 0.5 to 36 m s*^−^*^1^. The competitors’ curves are single-peaked and the peaks are differently placed: SDLfilter at 1.0, atlastools and the naïve cap at 0.75, trip::sda broadly flat to 5. None reaches the parameter-free cascade at any ceiling. Because each optimum can be located only by scoring against a clean reference, a user without one cannot reach even these values except by chance.

**Table 11:** Mean *F*_1_ over the five contaminated tracks as a function of the speed ceiling supplied, in m s*^−^*^1^. The parameter-free cascade takes no ceiling and is constant by con- struction. The two intermediate rows give the cascade a ceiling it does not require, under each of the two available speed-peel modes.

| Tool | 0.5 | 0.75 | 1 | 1.5 | 3 | 5 | 10 | 20 | 36 |
| --- | --- | --- | --- | --- | --- | --- | --- | --- | --- |
| **cascade** (no parameter) | 0.865 | 0.865 | 0.865 | 0.865 | 0.865 | 0.865 | 0.865 | 0.865 | 0.865 |
| cascade + $v_{\max}$, asymmetric peel | 0.754 | 0.887 | 0.887 | 0.887 | 0.887 | 0.887 | 0.887 | 0.907 | 0.898 |
| cascade + $v_{\max}$, default peel | 0.473 | 0.569 | 0.569 | 0.597 | 0.604 | 0.643 | 0.643 | 0.757 | 0.852 |
| **SDLfilter** | 0.734 | 0.822 | 0.830 | 0.804 | 0.755 | 0.718 | 0.609 | 0.499 | 0.387 |
| **trip::sda** | 0.736 | 0.736 | 0.736 | 0.736 | 0.736 | 0.736 | 0.723 | 0.719 | 0.716 |
| **atlastools** | 0.723 | 0.795 | 0.795 | 0.724 | 0.719 | 0.576 | 0.574 | 0.313 | 0.158 |
| naive cap | 0.490 | 0.581 | 0.581 | 0.551 | 0.546 | 0.448 | 0.446 | 0.281 | 0.165 |

The two intermediate rows require comment, because the effect of giving the cascade a ceiling depends entirely on which pre-peel runs, and reporting either row alone would misattribute the result. Supplying v_max runs an iterative speed peel ahead of the detectors. Its default, pre_peel_aux = "none", is symmetric: a step exceeding the ceiling identifies an offending edge but not which of its two endpoints is at fault, so both are removed, and a one-location spike therefore costs its two legitimate neighbours as well. The optional "detectors" mode scores both endpoints from a single bridge-and-detour pass and removes only the worse one.

True positives are invariant at 141 across every ceiling and both modes, exactly the number the parameter-free cascade recovers. Recall is already saturated, so a ceiling cannot add a detection and can only change the count of legitimate locations removed with the outliers: from 39 under the parameter-free default to 310 under the symmetric peel at 0.5 m s*^−^*^1^, or to roughly half the parameter-free figure under the asymmetric peel.

The asymmetric mode therefore scores above the parameter-free cascade on this cohort, at a mean *F*_1_ of 0.887 to 0.907 against 0.865, and is nearly indifferent to the ceiling over a 48-fold range from 0.75 to 36 m s*^−^*^1^. I report this as a measurement and draw no recommendation from it. The cohort is a single stochastic family with injected outliers of known geometry, and a setting selected because it maximises a score on such data is selected for the generator rather than for the animal. The package’s symmetric default was chosen on evidence this cohort cannot supply: its audit record reports the asymmetric peel winning on these synthetic tracks and regressing on a real one. Where a user has both a trustworthy ceiling and predominantly single-location spikes, the asymmetric mode is available and documented; the default recommendation of this paper remains the parameter-free cascade.

### S5.2 Irregular sampling

The cohort is irregular on four of its six tracks and exactly regular on the other two: the gap coefficient of variation is 1.12, 1.02, 0.96 and 1.13 on CPF_A–D, whose schedules are duty-cycled with dropouts, and 0 on CPF_E and CPF_F. What the cohort does not provide is a *controlled* contrast, since irregularity and sample size vary together across those tracks. Table 12 re-runs the comparison under three thinned schedules. Ground-truth indices are re-derived after thinning, so an injected outlier removed by the schedule is not counted against any tool.

**Table 12:** Mean *F*_1_ under four sampling schedules: the cohort as published; every second location retained, which halves the data while leaving the spacing regular; a day–night duty cycle; and that cycle interrupted by clustered acquisition failures. The thinned column is the control that separates sample size from gap structure. Every column averages the same three tracks, CPF_A, CPF_D and CPF_E: halving the data removes every injected outlier from CPF_C and CPF_F, leaving *F*_1_ undefined there, so a common set is the only like-for-like comparison. The regular column is therefore not the five-track mean reported in Section 3.

| Tool | regular | 50% thinned | duty cycle | duty + dropouts |
| --- | --- | --- | --- | --- |
| **cascade** | 0.866 | 0.820 | 0.848 | 0.863 |
| **SDLfilter** | 0.710 | 0.704 | 0.677 | 0.677 |
| **trip::sda** | 0.671 | 0.708 | 0.695 | 0.690 |
| **atlastools** | 0.651 | 0.662 | 0.648 | 0.629 |
| naive cap | 0.530 | 0.577 | 0.560 | 0.557 |

Two schedules retain a comparable share of the locations but differ in gap structure, and the regular-thinning column is the control. Its interpretation is the point of the design: the cascade loses 0.046 in mean *F*_1_ when the data are halved with the spacing left regular, and 0.003 under the duty cycle with dropouts, at 59% retention and a gap coefficient of variation of 1.24. Reduced sample size, not gap irregularity, is what degrades its detection. The competitors do not share that pattern, and the comparison does not favour the cascade: SDLfilter loses 0.007 to the same halving, while trip::sda and atlastools score higher on the thinned data than on the full track. Of the tools tested, the cascade is the one most sensitive to having less data. One qualification: the duty-cycle schedule retains more locations than the regular thinning, so that particular pair is not a clean contrast; the comparison that holds is the duty cycle with dropouts against regular sampling.

### S5.3 Positional error

The generator places legitimate locations exactly on the latent path, so recall of 1.000 and zero false positives on the clean reference are properties of a noise-free simulation. Table 13 adds heteroscedastic per-location error, 3% of locations drawn at five times the nominal standard deviation, to every location after outlier injection, and runs all five tools at each level. The cascade is not told the error magnitude, since the claim under test is that it requires no such parameter.

**Table 13:** Mean *F*_1_ over the five contaminated tracks against per-location positional error, in metres. Level 0 reproduces the published cohort. The 100 m column is retained for completeness and discussed in the text.

| Tool | 0 | 10 | 30 | 100 |
| --- | --- | --- | --- | --- |
| **cascade** | 0.865 | 0.906 | 0.846 | 0.692 |
| **SDLfilter** | 0.804 | 0.800 | 0.794 | 0.504 |
| **trip::sda** | 0.736 | 0.736 | 0.736 | 0.735 |
| **atlastools** | 0.724 | 0.724 | 0.724 | 0.694 |
| naive cap | 0.551 | 0.551 | 0.551 | 0.491 |

The cascade leads at 10 and 30 m, and its mean *F*_1_ at 10 m exceeds its noise-free value, which is consistent with a small amount of scatter separating the score distributions that the detectors threshold. At 100 m it falls to 0.692 and trip::sda, which is almost insensitive to error at any level, leads at 0.735. I retain that column rather than truncate the range, but it describes a regime outside the method’s intended input: a 100 m standard deviation per location at a 15-minute interval is not scatter about a good position but a population of cold-start acquisitions, which are identified from fix geometry rather than from trajectory shape and are removed by mt_filter_gps_quality() as recommended in Section 4. A screening cascade applied to unfiltered data of that quality is being asked to do a job the tag metadata already answers.

### S5.4 Worst-case behaviour across the cohort

Mean *F*_1_ across tracks, the summary Section 3 leads with, is a poor description of a screening tool: it lets a method that is excellent on four tracks and useless on the fifth look competent, and a user cannot know in advance which track they hold, establishing that is what the cleaning step is for. Table 14 therefore reports each tool’s *worst* track as well as its average, at two levels of per-location positional error, over one hundred replicate realisations.

**Table 14:** Worst-case and mean detection skill per tool across the five contaminated tracks, over one hundred replicate realisations, at two levels of per-location positional error. *Worst track* is the track on which that tool scores lowest. The competitors each hold *v*_max_ = 1.5 m s*^−^*^1^; the cascade holds nothing. Generated by scripts/robustness_summary.R.

| Error | Tool | Worst track | Worst $F_1$ | Worst recall | Mean $F_1$ |
| --- | --- | --- | --- | --- | --- |
| 0 m | **cascade** (no parameter) | CPF_D | 0.638 | 0.973 | 0.884 |
|  | SDLfilter | CPF_D | 0.270 | 0.256 | 0.803 |
|  | naive cap | CPF_D | 0.062 | 0.033 | 0.512 |
|  | trip::sda | CPF_D | 0.065 | 0.033 | 0.733 |
|  | atlastools | CPF_D | 0.000 | 0.000 | 0.656 |
| 30 m | **cascade** (no parameter) | CPF_F | 0.616 | 0.958 | 0.821 |
|  | SDLfilter | CPF_D | 0.196 | 0.160 | 0.777 |
|  | naive cap | CPF_D | 0.062 | 0.033 | 0.512 |
|  | trip::sda | CPF_D | 0.065 | 0.033 | 0.733 |
|  | atlastools | CPF_D | 0.000 | 0.000 | 0.656 |

Worst-case recall is the quantity to read. Unlike *F*_1_ it does not depend on the size of the truth set, so it is not inflated on a track carrying eighty injected outliers nor deflated on one carrying four: a distortion that matters here, because the same handful of false positives costs the cascade 0.03 of *F*_1_ on the former and 0.19 on the latter. And a screening step that misses three-quarters of the outliers on some deployments is unusable whatever its average.

All four competitors have their worst case on the coherent block, and all four fall to 0.256 or below there; the cascade’s minimum over the cohort is 0.973. The ordering is unchanged when per-location error is added, though the cascade’s own worst track then becomes the multi-state one. Two caveats belong with the table. The competitors are scored on a ceiling calibrated against a clean reference, so these are their best case, not their typical one. And, as noted above, the replicate cohort reproduces four of the six bundled tracks exactly and rebuilds the other two duty-cycled, so the multi-state row describes a related construction rather than the published track.

### S5.5 Block recovery against block size and position

The cohort contains one coherent block: 30 locations, mid-track, recovered in full. Table 15 asks how far that generalises, by holding an injected block’s construction fixed (the clean reference CPF_B as host, one seed, a *∼*120 km offset, the same tight within-block jitter), and varying only its size and its start position. The host flags nothing when untouched, so every flag counted is a response to the block.

Two things follow. At 30 locations, the cohort’s size, recovery is complete at every position tested, which is why a single cohort result reads as a general one. Away from that size it is not: recovery ranges from 1.000 down to 0.043, and a 300-location block at 85% of the way along the track is recovered 13 locations in 300. Where the expansion step does fire it recovers the block but can condemn a legitimate segment with it; the largest false-positive count in the sweep is 541.

The two extreme columns are the two true boundaries, and they are not interchange- able. At the trailing boundary, the block ending on the track’s last location, every size is recovered in full; at the leading boundary a 150-location block is recovered 105 locations in 150. The trailing column is also the configuration of the boundary-spoof demonstration in Supplement S4.2, and reproduces it exactly at 150 locations: 150 recovered, five false positives, *F*_1_ = 0.984, of which 146 carry the *block* class. The two tables therefore describe the same object where they overlap.

The percentages belong to this construction (one host, one offset, one jitter scale), and nothing here selects a configuration or tunes to the outcome. What the sweep establishes is the shape: recovery of a coherent block degrades with its size, the cohort samples the most favourable point on that surface, and a partially recovered block leaves a track that looks cleaned and is not.

## S6 Computational performance

The cascade is single-threaded per track and scans each detector linearly in track length up to a sort of its score vector, *O*(*n* log *n*); multi-individual datasets parallelise trivially across individuals. Wall-time scaling on synthetic tracks of geometrically increasing length is in Figure 5. The figure isolates track length on lightly contaminated tracks, and so measures the regime in which the pass count stays small; it cannot show the cost driver named above, which is contamination rather than length. The direction of that driver is visible on the cohort’s spoof track, where supplying a physiological cap shortens the run substantially: the pre-peel removes the impossible locations in one convergent pass instead of leaving them to be peeled away over many. No figure is put on that here, for the reason given in the next sentence. The same cap costs recall on that track (1.000 *→* 0.900, Supplement S4.2), so a user choosing it is trading detection for time rather than getting both. Per-track timings are deliberately not tabulated: on this cohort they vary severalfold between runs of identical code with machine load, so a table of them would report the machine rather than the method.

**Figure 5:**
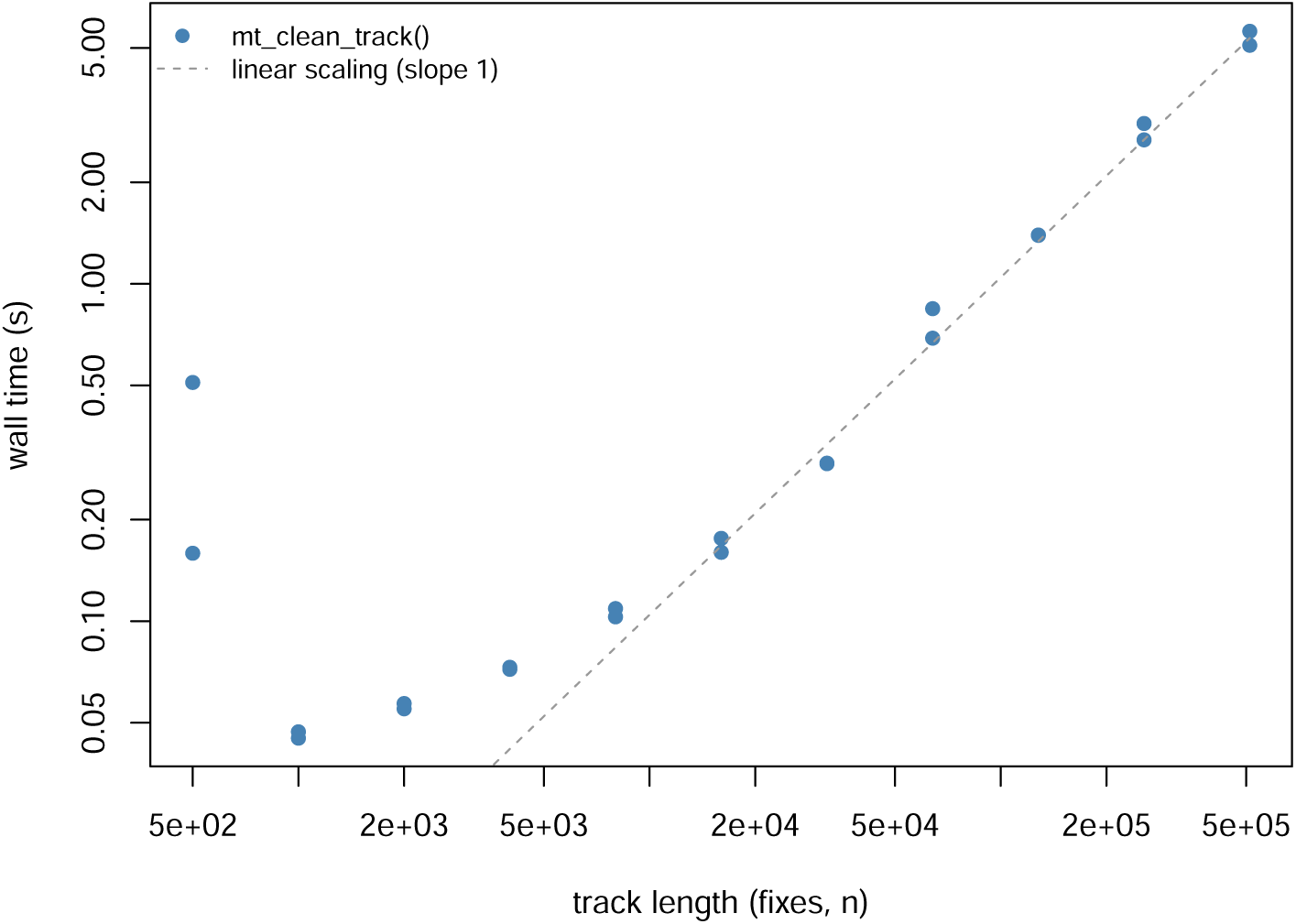
Wall-time scaling of mt_clean_track() on synthetic OUF tracks of geometrically increasing length, from 500 to 512,000 locations (two replicates per size). The dashed line marks ideal linear ( (*n*)) scaling, anchored in the asymptotic regime; fixed per-call overhead dominates below a few thousand locations. Generated by scripts/runtime_scaling.R.

## S7 Synthetic cohort: per-class flag composition and confusion matrices

Table 17 reports, for each cohort track under the parameter-free default, the detection confusion summary (true and false positives, false negatives and *F*_1_) alongside the composition of the flags across the four error_class values that arise on this cohort (the numeric backing of Figure 6). Every track recovers all injected outliers (false negatives are zero throughout); the false positives are the conservative over-flagging discussed in Section 3.1. The class columns may sum to fewer than the flagged total: a small number of flags are raised on cascade paths that do not assign an error_class label, and are not tabulated by class here. The block and physiological classes do not appear, as the cohort exercises neither block expansion (Supplement S4.2) nor a supplied speed cap at default settings.

**Figure 6:**
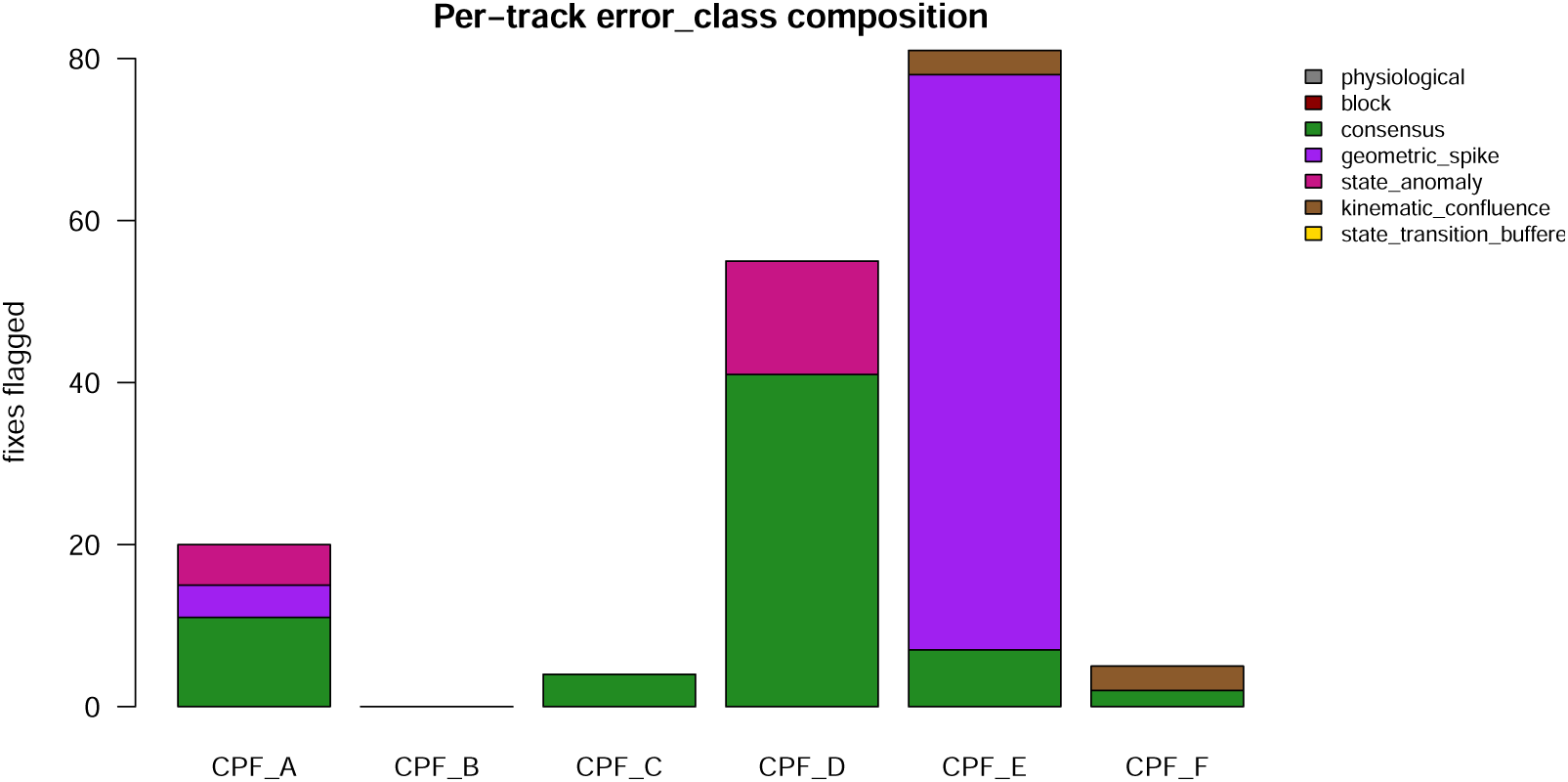
Per-track. error_class **composition across the synthetic cohort** under the parameter-free default. Stacked counts of the flag classes that arise on this co- hort: consensus, geometric_spike, state_anomaly, and kinematic_confluence. Two further classes of the taxonomy are not exercised here at default settings: block re- quires the graph-based block-expansion step (demonstrated on the boundary-spoof track of Supplement S4), and physiological requires a user-supplied speed cap. The composition is itself diagnostic: a track dominated by geometric_spike indicates a colony-halo error structure (CPF_E); a spoof block recovered by consensus and state_anomaly (CPF_D); isolated spikes by consensus with a few geometric_spike/state_anomaly (CPF_A, CPF_C). See scripts/worked_examples. R.

### S7.1 Decision-rule comparison and sensitivity to construction

This cohort does not separate the default evidence_corroborated rule from the class_aware conjunction, and was not built to: no experiment here scores both. That the two should be close is expected rather than coincidental: both draw on the same four detectors and differ only in how they pool them, so once the detectors agree on the easy cases and fall silent on the clean ones, most of any track, the pooling rule has little left to decide. evidence_corroborated is the default not because it detects more but because, at comparable skill, it is the better-behaved object: it returns a continuous combined_evidence score rather than a bare flag, so a user can rank locations by suspicion or set their own operating point, and it extends to a new detector without rewriting a logical table.

### S7.2 Replicate-realisation confidence intervals

The cohort table in Section 3.1 (Table 16) reports a single realisation of each track. Because detection skill varies with the realisation seed, I regenerated the cohort across one hundred seeds at a fixed construction setting (nominal outlier magnitude, 30 m per-location positional error) and scored every tool on each realisation. What the regeneration reproduces needs stating precisely, because it is not the whole cohort: CPF_A, CPF_B, CPF_C and CPF_D come back exactly (same location counts, same truth-set sizes), but CPF_E and CPF_F are rebuilt duty-cycled, at 790 locations against the bundled 1,440 and 1,666 against 2,880, where the bundled tracks are regularly sampled. Results for those two tracks therefore describe a related construction rather than the published track, and are not replicates of it. Table 18 reports the per-track mean and 95% confidence interval of recall, precision, and *F*_1_ over the one hundred realisations. Recall stays high on every track but is not invariably 1 (CPF_A averages 0.958, the lowest in the cohort), and precision, the discriminating axis, is stable on the easy tracks and low on the spoof block (CPF_D, *≈* 0.48) as in the single realisation. CPF_F’s replicate mean *F*_1_ is 0.616 (95% CI 0.576– 0.656), below the single-realisation 0.73 of Table 16; the two are not directly comparable, since this track is among the two rebuilt on a different sampling schedule. That a single draw is an unreliable point estimate is shown by the confidence intervals themselves rather than by any one such comparison. On the clean reference, which carries no truth set, the cascade raised one false positive across the hundred realisations. Generated by scripts/replicate_ci.R.

**Table 15:** Fraction of an injected coherent block recovered by the parameter-free cascade, by *any* route, against block size (rows) and the block’s start position as a fraction of track length (columns). The block’s construction is identical in every cell. Generated by scripts/block_sweep.R

| Block size | 0% | 5% | 15% | 30% | 50% | 70% | 85% | 95% | 100% |
| --- | --- | --- | --- | --- | --- | --- | --- | --- | --- |
| 15 fixes | 1.00 | 1.00 | 1.00 | 0.67 | 1.00 | 1.00 | 1.00 | 1.00 | 1.00 |
| 30 fixes | 1.00 | 1.00 | 1.00 | 1.00 | 1.00 | 1.00 | 1.00 | 1.00 | 1.00 |
| 60 fixes | 1.00 | 1.00 | 1.00 | 0.27 | 1.00 | 0.33 | 1.00 | 1.00 | 1.00 |
| 150 fixes | 0.70 | 1.00 | 1.00 | 0.35 | 0.68 | 0.68 | 1.00 | 0.63 | 1.00 |
| 300 fixes | 1.00 | 1.00 | 0.36 | 0.37 | 0.37 | 0.29 | 0.04 | 1.00 | 1.00 |

**Table 16:** Per-track detection metrics for the zero-parameter cascade across the syn- thetic cohort. n is location count; truth is the size of the bundled ground-truth outlier set; flagged/TP/FP/FN are the locations flagged and the true-positive, false-positive, and false-negative counts against that truth set; *F*_1_ is the harmonic mean of precision and recall (1 = perfect). Auto-generated by scripts/worked_examples.R into figures/synth_metrics.tex.

| Track | $n$ | truth | flagged | TP | FP | FN | recall | precision | $F_1$ |
| --- | --- | --- | --- | --- | --- | --- | --- | --- | --- |
| CPF_A | 1748 | 23 | 25 | 23 | 2 | 0 | 1.00 | 0.92 | 0.96 |
| CPF_B | 3537 | 0 | 0 | 0 | 0 | 0 | NA | NA | NA |
| CPF_C | 185 | 4 | 4 | 4 | 0 | 0 | 1.00 | 1.00 | 1.00 |
| CPF_D | 1699 | 30 | 63 | 30 | 33 | 0 | 1.00 | 0.48 | 0.65 |
| CPF_E | 1440 | 80 | 81 | 80 | 1 | 0 | 1.00 | 0.99 | 0.99 |
| CPF_F | 2880 | 4 | 7 | 4 | 3 | 0 | 1.00 | 0.57 | 0.73 |

**Table 17:** Per-track detection summary and error_class composition under the zeroparameter cascade. The confusion columns (TP, FP, FN, F1) are as in Table 16; the remaining columns count the locations in each flag class, abbreviated cons. (consensus), g.spike (geometric_spike), s.anom (state_anomaly), k.confl (kinematic_confluence). Unclas. counts flagged locations that satisfy no class predicate and so carry error_class = NA: the default rule decides on summed evidence, which a detector can contribute without firing, so a location can clear it while fewer than two detectors fire and no class predicate matches; all 15 such locations in the cohort have exactly one detector firing, so the class columns need this term to sum to flagged. Wall time is not reported: it varies severalfold between runs of identical code with machine load, and the scaling claim rests on Figure 5 rather than on per-track timings. From figures/synth_metrics.csv, figures/synth_class_composition.csv and scripts/worked_examples.R.

| Track | Detection |  |  |  | Flag class |  |  |  |  |  |
| --- | --- | --- | --- | --- | --- | --- | --- | --- | --- | --- |
| | TP | FP | FN | $F_1$ | cons. | g.spike | s.anom | k.confl | unclas. | flagged |
| CPF_A | 23 | 2 | 0 | 0.958 | 11 | 4 | 5 | 0 | 5 | 25 |
| CPF_B | 0 | 0 | 0 | — | 0 | 0 | 0 | 0 | 0 | 0 |
| CPF_C | 4 | 0 | 0 | 1.000 | 4 | 0 | 0 | 0 | 0 | 4 |
| CPF_D | 30 | 33 | 0 | 0.645 | 41 | 0 | 14 | 0 | 8 | 63 |
| CPF_E | 80 | 1 | 0 | 0.994 | 7 | 71 | 0 | 3 | 0 | 81 |
| CPF_F | 4 | 3 | 0 | 0.727 | 2 | 0 | 0 | 3 | 2 | 7 |

**Table 18:** Replicate-realisation per-track skill of the zero-parameter cascade: mean and 95% confidence interval of recall, precision, and *F*_1_ over one hundred cohort realisations at a fixed construction setting, with 30 m per-location positional error. CPF_B is the clean track (no truth set). CIs are clamped to [0, 1]. From scripts/replicate_ci.R via figures/synth_class_composition.csv and scripts/worked_examples.R.

| Track | recall | | precision | | $F_1$ |
| --- | --- | --- | --- | --- | --- |
|  | mean | [95% CI] | mean | [95% CI] | mean [95% CI] |
| CPF_A | 0.96 | [0.95, 0.96] | 0.95 | [0.93, 0.96] | 0.95 [0.93, 0.96] |
| CPF_B | — | — | — | — | — |
| CPF_C | 0.97 | [0.96, 0.99] | 0.93 | [0.89, 0.98] | 0.93 [0.89, 0.97] |
| CPF_D | 1.00 | [0.99, 1.00] | 0.49 | [0.48, 0.49] | 0.65 [0.65, 0.66] |
| CPF_E | 1.00 | [1.00, 1.00] | 0.92 | [0.92, 0.93] | 0.96 [0.96, 0.96] |
| CPF_F | 1.00 | [0.99, 1.00] | 0.48 | [0.43, 0.53] | 0.62 [0.58, 0.66] |

## Notes

https://github.com/move2universe/move2utils

### Competing Interest Statement

The authors have declared no competing interest.

### Summary of Updates

Tightened text, updated version numbers of the move2utils package. Rewrote abstract

